# Shared and distinct functional effects of patient-specific *Tbr1* mutations on cortical development

**DOI:** 10.1101/2022.01.27.478064

**Authors:** Marissa Co, Rebecca A. Barnard, Jennifer N. Jahncke, Sally Grindstaff, Lev M. Fedorov, Andrew C. Adey, Kevin M. Wright, Brian J. O’Roak

## Abstract

Predicted loss-of-function and missense heterozygous *de novo* mutations of *TBR1* are strongly associated with intellectual disability and autism. The functional effects of these heterogeneous mutations on cortical development and genotype-phenotype relationships have yet to be explored. We characterized mouse models carrying patient mutations A136PfsX80 and K228E, finding convergent and discordant phenotypes. The A136PfsX80 mutation is loss-of-function and allelic to the *Tbr1* knockout. In contrast, K228E causes significant upregulation of TBR1. Heterozygosity of either mutation produces axon defects, including reduction of the anterior commissure, and CTIP2 downregulation in adult cortex. While mice lacking TBR1 show extensive cortical apoptosis and inverted layering, K228E homozygotes show normal apoptosis levels and a complex layering phenotype—suggesting partial, yet abnormal, function of the allele. The construct and face validity of these *Tbr1* patient mutation mice suggests they will be valuable translational models for studying the function of this essential brain transcription factor.

## INTRODUCTION

Neurodevelopmental disorders (NDDs) impact numerous individuals and their caretakers worldwide: frequencies for autism spectrum disorder (ASD) approach 1%, intellectual disability (ID) 1%, and attention-deficit/hyperactivity disorder (ADHD) 2.5%–5% (American Psychiatric Association, 2013). These conditions are highly heritable, yet their genetic architectures are complex, with ASD risk factors ranging from monogenic disruptions to polygenic summations, or with variable expressive loci contributing to multiple neurodevelopmental phenotypes (Iakoucheva et al., 2019; Lee et al., 2019). A considerable source of NDD genetic risk is *de novo* mutations, which account for 30–50% of ASD, ID, and/or broader developmental disorder cases (Hamdan et al., 2014; Iossifov et al., 2014; McRae et al., 2017; Yoon et al., 2021). Such mutations are informative for dissecting biological mechanisms underlying NDDs, as they typically induce large phenotypic effect sizes and impair single genes.

In the past decade, *TBR1* (T-Box Brain Transcription Factor 1) has emerged as a critical gene for understanding NDD risk via *de novo* mutations. *TBR1* was among the first ASD candidate genes identified via exome sequencing of families with sporadic ASD, and was subsequently validated as a high-confidence risk gene through targeted resequencing (Neale et al., 2012; O’Roak et al., 2014; O’Roak et al., 2012a; O’Roak et al., 2012b). Over 100 risk variants impacting *TBR1* have been reported in the ClinVar database, and genetic constraint metrics indicate that *TBR1* is highly intolerant to both loss-of-function and missense mutations (Karczewski et al., 2020; Landrum et al., 2018). Accordingly, no biallelic *TBR1* mutations have been identified in humans, and *Tbr1* knockout mice die perinatally (Bulfone et al., 1998; Nambot et al., 2020). *TBR1*’s spatiotemporal expression in developing human brain places it at the center of an ASD risk gene co-expression network specific to mid-fetal glutamatergic cortical neurons (Willsey et al., 2013). During the equivalent period in mouse cortex, TBR1 directly binds and regulates the expression of other high-confidence ASD risk genes (Notwell et al., 2016). *Tbr1* is expressed in several early-born neuronal populations essential for proper mouse corticogenesis, including subplate neurons, Cajal-Retzius cells, and deep-layer glutamatergic neurons, and its complete knockout consequently impairs cortical layer formation, cell survival, neuronal fate acquisition, and axon tract formation (Bedogni et al., 2010; Han et al., 2011; Hevner et al., 2001; McKenna et al., 2011). Thus, *TBR1* may play a major role in NDD risk by virtue of its coexpression with and regulation of other high-confidence ASD genes during embryonic cortical development.

Individuals with *de novo TBR1* mutations exhibit moderate-to-severe ID and/or developmental delay, ASD or autistic traits, and behavior disorders such as attention-deficit and aggression (McDermott et al., 2018; Nambot et al., 2020). Over half of patients examined by MRI showed anterior commissure reduction, hippocampal dysplasia, and/or cortical dysplasia with gyral anomalies (Nambot et al., 2020; Vegas et al., 2018). Likewise, heterozygous *Tbr1* knockout mice (*Tbr1^+/–^*), which model human *TBR1* haploinsufficiency, show reduction of the anterior commissure as well as cognitive and social deficits (Huang et al., 2014). Mice in which *Tbr1* is conditionally deleted (*Tbr1^cKO^*) from deep-layer cortical glutamatergic neuronal subpopulations also show NDD-relevant behaviors, such as social deficits and aggression (Fazel Darbandi et al., 2020; Fazel Darbandi et al., 2018). While convergent defects in dendritic spine formation were seen in both *Tbr1^+/–^* and *Tbr1^cKO^* mice, the deletion of *Tbr1* several days after its initial expression in *Tbr1^cKO^* models may limit their translatability. Moreover, *Tbr1* deletion models do not capture the allelic heterogeneity of human *de novo TBR1* mutations. *In vitro* studies have shown that different *TBR1* mutations produce mutant proteins with varying stability, transcriptional activity, subcellular localization, and cofactor binding (den Hoed et al., 2018; Deriziotis et al., 2014). Furthermore, aside from their cognitive and autistic traits, *TBR1* patients exhibit considerable phenotypic heterogeneity in other symptoms, such as motor impairment, brain malformations, and abnormal EEG, and whether these can be attributed to unique genotype-phenotype relationships is currently unknown (Nambot et al., 2020).

Mice harboring the K228E patient mutation in the T-box DNA-binding domain of TBR1 were recently generated and characterized (Yook et al., 2019). In contrast to deletion mutants, *Tbr1^+/K228E^* mice showed elevated TBR1 protein levels, and the mutant TBR1-K228E protein exhibited decreased affinity for DNA and increased stability *in vitro*. While transcriptional dysregulation was observed in embryonic forebrains of *Tbr1^+/K228E^* and *Tbr1^K228E/K228E^* mice, the overall *in vivo* functionality of the mutant allele remains unclear, as do its effects on aspects of neurodevelopment such as axon tract formation. Aside from K228E, no other *TBR1* patient mutations have been modeled *in vivo*.

Here, we generated two patient-specific *Tbr1* mutant mouse lines carrying an early truncating frameshift (A136PfsX80) or a missense mutation (K228E) in order to ascertain the *in vivo* functional effects of these mutations. We also characterized an in-frame *Tbr1* deletion (Δ348– 353) encompassing five reported human *TBR1* variants. With the *Tbr1* knockout mouse line as a comparison (Bulfone et al., 1998), we used molecular, histological, and genetic approaches to determine the impacts of these mutations on *Tbr1* expression and cortical development. The following results provide new insights into the regulation and function of a critical NDD risk gene during cortical development and allow for enhanced *in vivo* modeling of *TBR1*-related disorders.

## RESULTS

### Generation of mouse lines modeling *Tbr1* patient mutations

Using CRISPR/Cas9 genome editing (Aida et al., 2015), we generated three *Tbr1* mouse lines: *Tbr1^A136PfsX80^* (c.402del; p.A136PfsX80), *Tbr1^K228E^* (c.682A>G; p.K228E), and *Tbr1^Δ348–353^* (c.1042_1059del; p.E348_P353del). The sequence changes producing these predicted mutant proteins are identical between human and mouse, and while c.402 is cytosine in human and thymine in mouse, their respective codons are synonymous. A136PfsX80 is a frameshift mutation predicted to yield a truncated protein missing the T-box DNA-binding domain, while K228E is a missense mutation within the T-box (Figure 1A) (O’Roak et al., 2012a; O’Roak et al., 2012b). Both mutations impact highly conserved residues among vertebrates, and the residues following the frameshift of A136PfsX80 are conserved between human and mouse (Figure 1B-C and Figure S1C). For these lines, we tested multiple CRISPR synthetic guide RNA (sgRNA) sequences and designed single-stranded oligo DNA donors (ssODNs) to knock-in the patient mutations through homology-directed repair (Table S1). The in-frame deletion mutant Δ348–353 was generated through chance non-homologous end joining during editing intended to generate a different point mutation. This deletion is also located within the T-box and encompasses five reported human *TBR1* variants in ClinVar and gnomAD: two nonsense mutations classified as pathogenic or likely pathogenic (S351X, Q352X), and three missense mutations of uncertain or conflicting significance (T350A, S351R, P353A) (Figure S1A-B, Table S2) (Karczewski et al., 2020; Landrum et al., 2018). Despite the strong human-mouse conservation of the T-box, residues 348–353 fall within a poorly conserved site among other mouse T-box family proteins (Figure S1C).

**Figure 1.**
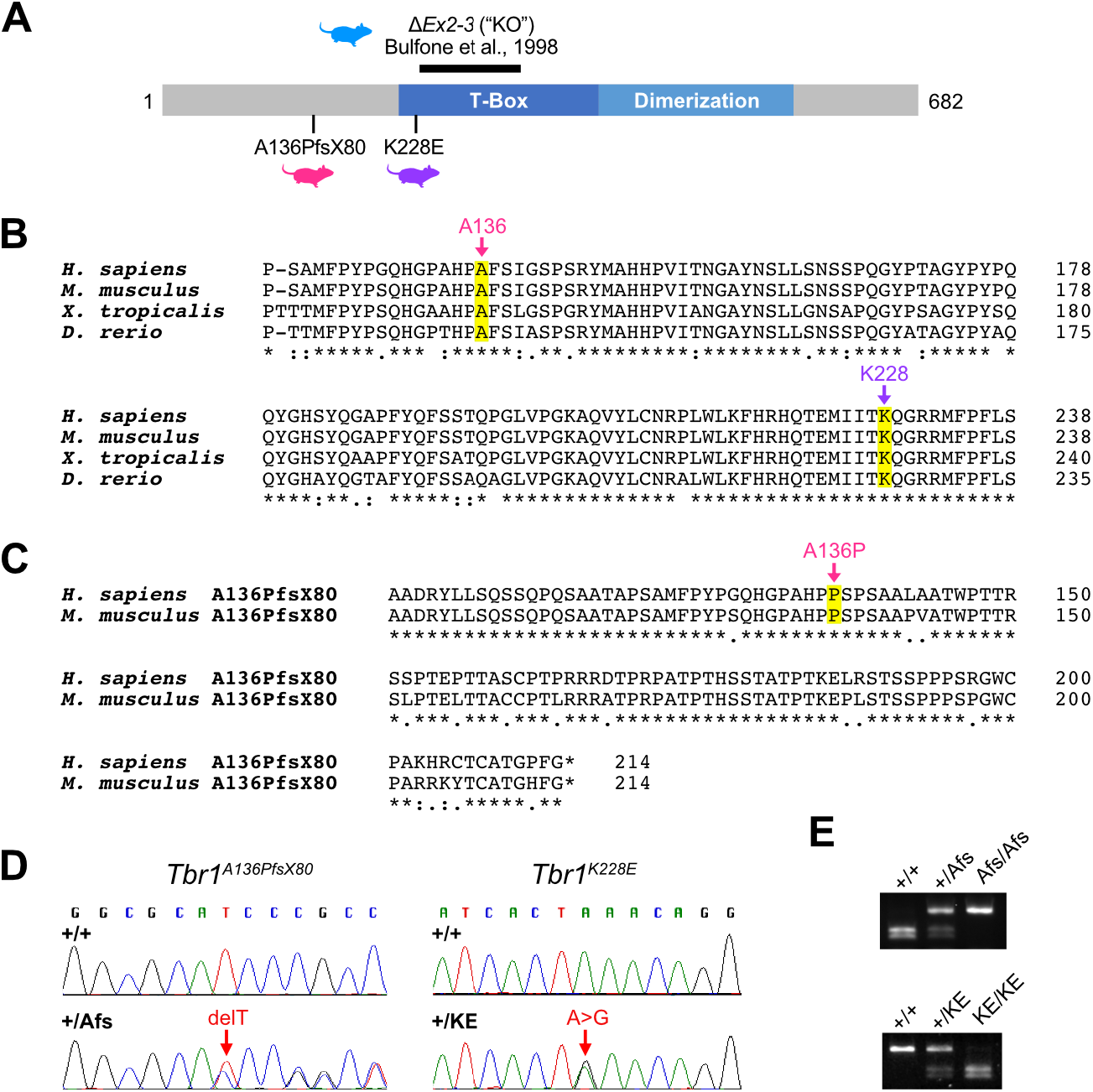
Generation of Mouse Lines Carrying *Tbr1* Patient Mutations. (A) Schematic of TBR1 protein summarizing mouse lines used in this study: published exon 2-3 knock-out line (Bulfone et al., 1998) and newly generated CRISPR knock-in lines carrying a frameshift (A136PfsX80) or missense (K228E) patient mutation. (B) Multiple sequence alignment of TBR1 A136 and K228 sites conserved across vertebrate species. (C) Pairwise alignment of predicted frameshift regions of human and mouse TBR1-A136PfsX80 protein. (D) Sanger sequencing of genomic DNA showing thymine (T) deletion in *Tbr1^A136PfsX80^* heterozygote (+/Afs) and A>G substitution in *Tbr1^K228E^* heterozygote (+/KE). (E) Enzyme digest-based PCR genotyping of *Tbr1* patient mutant mouse lines.

Using Sanger sequencing, we confirmed the *Tbr1* mutations and lack of local region (~300 bp) off-target edits in the patient mutant line founders (F0) and their offspring (Figure 1D). The sgRNAs used had limited potential for exonic off-target effects (≥2 off-target mismatches). However, to account for any off-target edits, we backcrossed each line to the parental C57BL/6NJ strain for at least two generations and always compared littermate controls and mutants within experiments. For *Tbr1^A136PfsX80^*, we also characterized three separate F1-descendant branches and identified no brain phenotypic differences among these lineages, further suggesting that any phenotypes observed were specific to the primary *Tbr1* editing event. After establishing these lines, patient mutant mice were genotyped via PCR with restriction enzyme digest (Figure 1E).

Heterozygous mutants from the *Tbr1^A136PfsX80^*, *Tbr1^K228E^*, and *Tbr1^Δ348–353^* lines (*Tbr1^+/Afs^*, *Tbr1^+/KE^*, and *Tbr1^+/Δ^*, respectively), as well as *Tbr1^Δ/Δ^* homozygotes, appeared healthy with normal outward morphology. In contrast, *Tbr1^Afs/Afs^* and *Tbr1^KE/KE^* mice died perinatally and had small olfactory bulbs, as was previously described for *Tbr1^−/−^* mice (Figure S2A) (Bulfone et al., 1998). Mutants from the *Tbr1^A136PfsX80^* and *Tbr1^K228E^* lines did not differ from wild-type (WT) control littermates in neonatal brain or body size, nor did heterozygous mutants show gross motor impairments across postnatal development (Figure S2B-D). We proceeded to characterize these lines in comparison with an established *Tbr1* knockout (*Tbr1^KO^*) line generated by replacement of exons 2 and 3 with a neomycin cassette and backcrossed to a C57BL/6NJ background (Figure 1A) (Bulfone et al., 1998).

### Contrasting effects of patient mutations on TBR1 protein levels in cortex

We first measured TBR1 protein levels in postnatal day (P) 0 cortex from *Tbr1^KO^* mice and the patient mutant mouse lines. Full-length TBR1 signal was reduced by approximately 40% in *Tbr1^+/–^* and *Tbr1^+/Afs^* and completely absent in *Tbr1^−/−^* and *Tbr1^Afs/Afs^* (Figures 2A-B). The predicted truncated TBR1-A136PfsX80 protein was also not detected using an N-terminal TBR1 antibody. In contrast, TBR1 levels were increased by 2-fold and 5-fold in *Tbr1^+/KE^* and *Tbr1^KE/KE^* cortex, respectively. These TBR1 alterations were maintained in adult heterozygous mutant cortex (Figure 2C). Despite the decreases in TBR1 protein in *Tbr1^KO^* and *Tbr1^A136PfsX80^* mutants, *Tbr1* transcript levels were not significantly different across genotypes, suggesting that the protein reductions resulted from post-transcriptional processes (Figure 2D). Notably, in P0 *Tbr1^+/Afs^* cortex, transcripts from the mutant allele comprised only a small proportion of overall *Tbr1* transcript, suggesting upregulation of WT transcript and/or nonsense-mediated decay of mutant transcript (Figure S3A-D). *Tbr1* transcript was increased by 1.5-fold and 3.5-fold in *Tbr1^+/KE^* and *Tbr1^KE/KE^*, respectively, indicating that their increased TBR1 protein levels can be attributed to transcriptional upregulation (Figure 2D). In *Tbr1^Δ348–353^* mutants, TBR1 protein and *Tbr1* transcript levels were similar to WT (Figure S2D-F).

**Figure 2.**
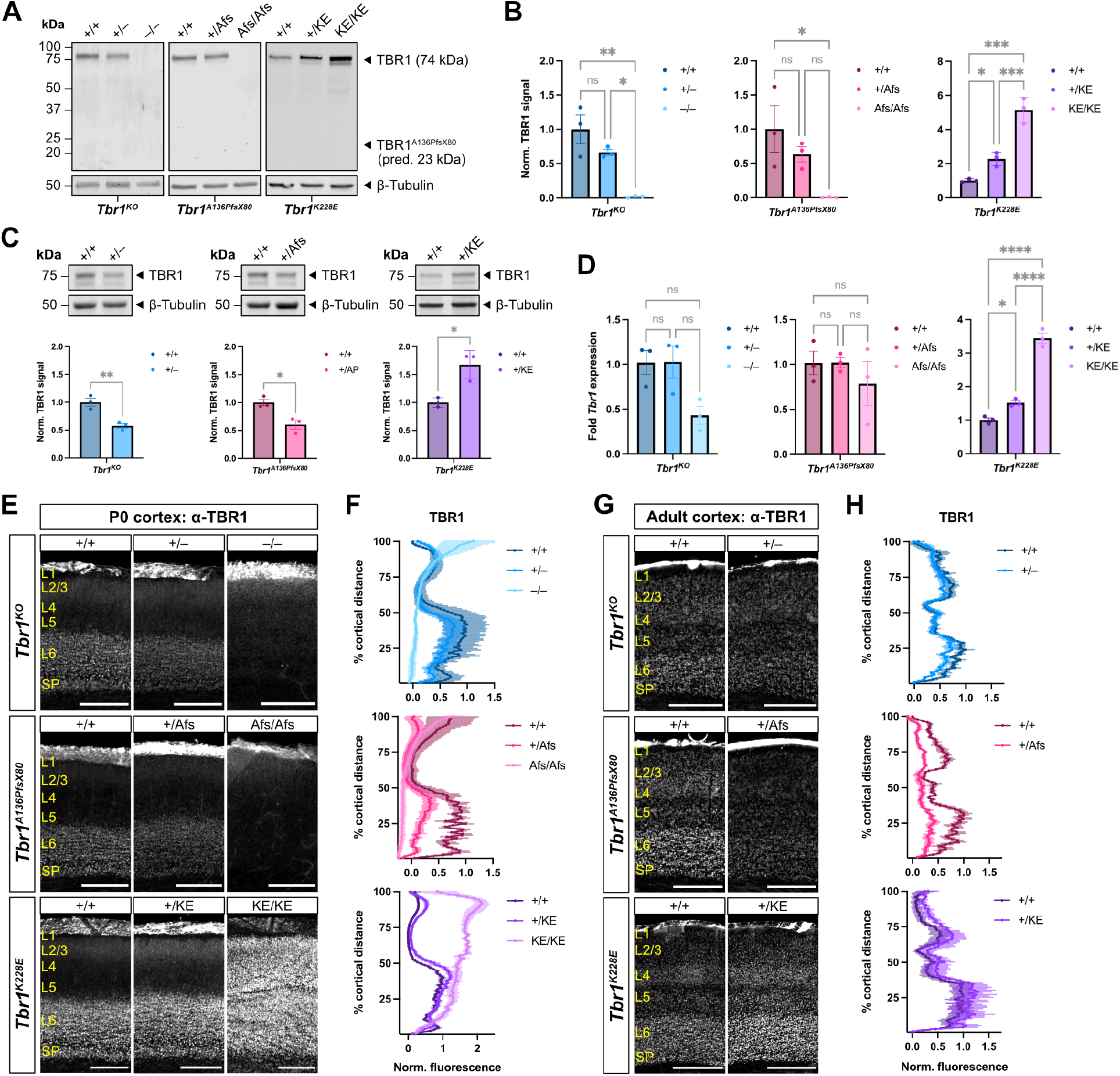
*Tbr1* A136PfsX80 is a Loss-of-Function Frameshift Mutation, while *Tbr1* K228E Missense Mutation Causes *Tbr1* Upregulation. (A-B) Western blots for TBR1 in postnatal day (P) 0 cortical lysates from *Tbr1* knock-out and patient lines (n = 3 mice per genotype). β-III-tubulin was used as loading control. Predicted molecular weight of TBR1 A136PfsX80 protein is indicated, but no truncated protein product is detected in *Tbr1^A136PfsX80^* mutants. (C) Western blots for TBR1 in adult cortical lysates from *Tbr1* knock-out and patient lines (n = 3 mice per genotype). (D) RT-qPCR for *Tbr1* in P0 *Tbr1* knock-out and patient line cortex (n = 3 mice per genotype). (E-H) TBR1 immunostaining and fluorescence intensity quantification in P0 and adult *Tbr1* knock-out and patient line somatosensory cortex (n = 3-4 mice per genotype). L: layer, SP: subplate. Scale bar = 200 μm in (E); 500 μm in (G). Data are plotted as mean ± SEM. Each dot represents one animal. One-way ANOVA with Tukey’s multiple comparisons test (B) and (D); unpaired Student’s t-test (C). *p < 0.05; **p < 0.01; ***p < 0.001; ****p < 0.0001; ns, not significant.

In the developing cortex, TBR1 is present at high levels in deep-layer excitatory projection neurons, and in adulthood is present in both upper- and deep-layer excitatory neurons (Hevner et al., 2001). To assess TBR1 layer distribution and levels at these stages, we immunostained for TBR1 in P0 and adult brain sections and plotted fluorescence profiles across the cortical layers. At both stages, heterozygotes of each mutant line showed TBR1 distributions similar to WT (Figure 2E-H). While *Tbr1^−/−^* and *Tbr1^Afs/Afs^* showed minimal TBR1 fluorescence at P0, *Tbr1^KE/KE^* unexpectedly showed substantial TBR1 upregulation across all cortical layers (Figure 2E-F). In these mice, the vast majority of NeuN+ cortical neurons expressed TBR1 (Figure S3E-F). In contrast to the patient lines, *Tbr1^Δ/Δ^* mice showed normal deep-layer TBR1 expression in P0 cortex (Figure S2G-H). Overall, these results show that A136PfsX80 causes absence of TBR1 protein, K228E causes elevated TBR1 levels due to transcriptional upregulation, and Δ348–353 has minimal impact on TBR1 levels.

### Homozygosity of *Tbr1* patient mutations A136PfsX80 and K228E causes distinct cortical layering defects

Cortical formation is an intricate process whereby newborn neurons migrate outward from a germinal zone to form six distinct cytoarchitectural layers organized by birthdate in an “inside-out” fashion (Kwan et al., 2012). Complete *Tbr1* knockout in mice causes *reeler*-like disorganization of cortical layering, fate-switch of layer (L) 6 neurons to L5 neurons, and abnormal distribution of cortical interneurons (Han et al., 2011; Hevner et al., 2001; McKenna et al., 2011). To examine cortical formation in the *Tbr1* patient mutant lines, we immunostained for layer markers CUX1 (L2-4) and CTIP2 (L5) at P0 and adulthood. At both stages, *Tbr1^+/–^*, *Tbr1^+/Afs^*, and *Tbr1^+/KE^* mice showed grossly normal layer formation and layer marker distributions in primary somatosensory (S1) cortex (Figure 3A-D). By adulthood, their cortical layering remained normal but they showed consistent reductions in CTIP2 fluorescence intensity in L6 (Figure 3C-D). We also immunostained for the major interneuron subclass markers somatostatin (SST) and parvalbumin (PV) in *Tbr1^+/–^*, *Tbr1^+/Afs^*, and *Tbr1^+/KE^* adult mice, but found no change in their density or distribution within S1 cortex (Figure S4). In contrast to heterozygous mutants, homozygous mutants showed major cortical layering defects differing by mutation (Figure 3A-B). Specifically, in *Tbr1^−/−^* and *Tbr1^Afs/Afs^* mice, layering appeared inverted, with CUX1+ cells mostly residing in inner cortex and CTIP2+ cells almost exclusive to outer cortex. *Tbr1^KE/KE^* mice instead showed a more complex layering phenotype, with CUX1+ cells forming a thinner mid-cortical layer and CTIP2+ cells residing in both outer and inner cortex. These abnormally formed layers were also revealed cytoarchitecturally by Hoechst nuclear staining. In each mutant line, homozygous mutants additionally showed an overabundance of CTIP2+ neurons compared to WT, suggesting L6→L5 neuronal fate-switching. These data indicate that heterozygosity of A136PfsX80 or K228E has minimal impacts on cortical layer formation but alters CTIP2 levels in mature L6, while homozygosity of these mutations has differential effects on cortical layer formation.

**Figure 3.**
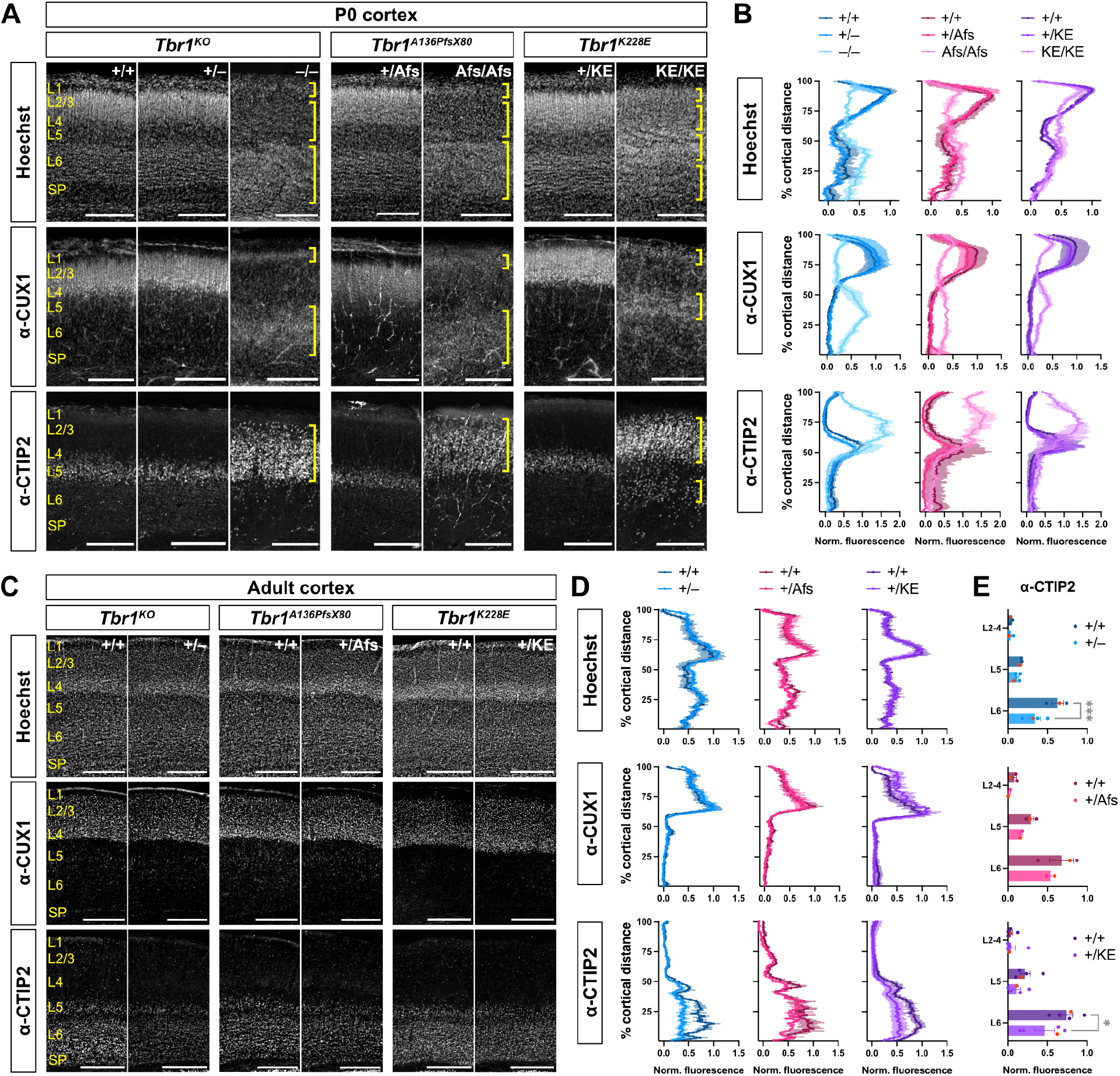
Effects of *Tbr1* Patient Mutations on Cortical Layer Formation and Layer Marker Expression. (A) Hoechst nuclear stain and immunostaining for cortical layer markers CUX1 (L2-4) and CTIP2 (L5) in postnatal day (P) 0 *Tbr1* knock-out and patient line somatosensory cortex. Yellow brackets indicate abnormal cortical layers formed in homozygotes. (B) Quantification of Hoechst, CUX1, and CTIP2 fluorescence intensity across the cortical mantle from (A) (n = 2-4 mice per genotype). (C-D) Hoechst, CUX1, and CTIP2 staining and quantification in adult *Tbr1* knock-out and patient line somatosensory cortex (n = 3-4 mice per genotype). (E) CTIP2 fluorescence binned by cortical layers using fluorescence values from (D). L: layer, SP: subplate. Scale bar = 200 μm in (A); 500 μm in (C). Data are plotted as mean ± SEM. Each dot represents one animal; red dots correspond to representative images. Two-way ANOVA with Šídák’s multiple comparisons test (E). *p < 0.05; ***p < 0.01.

To further assess the functionality of the K228E allele, we performed genetic complementation tests by crossing the *Tbr1^K228E^* line with the *Tbr1^KO^* line. *Tbr1^KE/–^* mice showed cortical layering defects partially reminiscent of *Tbr1^KE/KE^*, with cortex-wide TBR1 distribution and CTIP2+ cells present in both inner and outer cortex (Figure S5A-B). However, CUX1 and Hoechst distributions in *Tbr1^KE/–^* mice were more similar to those in *Tbr1^−/−^* mice. This result suggests that K228E over null is phenotypically intermediate between the two homozygous states, and confirms that the TBR1-K228E mutant protein has insufficient functionality to mediate proper cortical layer formation.

### T-box residues 348 to 353 are dispensable for normal TBR1 function during cortical formation

In contrast to homozygotes of the *Tbr1^KO^*, *Tbr1^A136PfsX80^*, *Tbr1^K228E^* lines, *Tbr1^Δ/Δ^* mice showed normal cortical layering and normal abundance of CTIP2+ neurons (Figure S2G-H). As with the *Tbr1^K228E^* line, we also assessed whether the Δ348–353 allele could complement the null allele by crossing this line to *Tbr1^KO^*. *Tbr1^Δ/–^* mice still showed normal TBR1 localization and layer formation, despite having only one copy of the mutant allele on a null background (Figure S5CD). We concluded that amino acids 348–353 of the T-box are dispensable for TBR1 function during cortical layer formation, and we limited further characterization of the *Tbr1^Δ348–353^* line to focus on the more pathogenic patient-specific mutations.

### *Tbr1* patient mutations A136PfsX80 and K228E cause equivalent axon tract defects

Previous studies identified requirement of *Tbr1* for normal axon tract development in the brain: *Tbr1^−/−^* mice show severe defects of the corpus callosum, anterior commissure, and internal capsule, while *Tbr1^+/–^* mice lack the posterior limb of the anterior commissure (Hevner et al., 2001; Huang et al., 2014). To examine axon tract formation in our *Tbr1* patient mutant lines, we immunostained for the axon markers L1 and neurofilament at P0 and adulthood, respectively. At these stages, the vast majority of heterozygotes from *Tbr1^KO^* and both patient mutant lines lacked an apparent posterior limb of the anterior commissure (n=11/11 total heterozygotes at P0; n=12/13 at adulthood), indicating a highly penetrant phenotype upon heterozygous *Tbr1* mutation (Figure 4A-B). One adult *Tbr1^+/KE^* mouse showed a very thinly formed posterior limb (Figure 4B, arrowhead). At P0, heterozygotes from all three lines also showed abnormal growth of external capsule axons. Homozygotes showed more severe axon defects, including disorganized and misdirected callosal fibers and complete absence of the anterior commissure (Figure 4A). We also examined the internal capsule, which contains corticothalamic and thalamocortical axon fibers, and observed grossly normal organization of this tract in heterozygotes and abnormal organization in homozygotes (Figure 4C).

**Figure 4.**
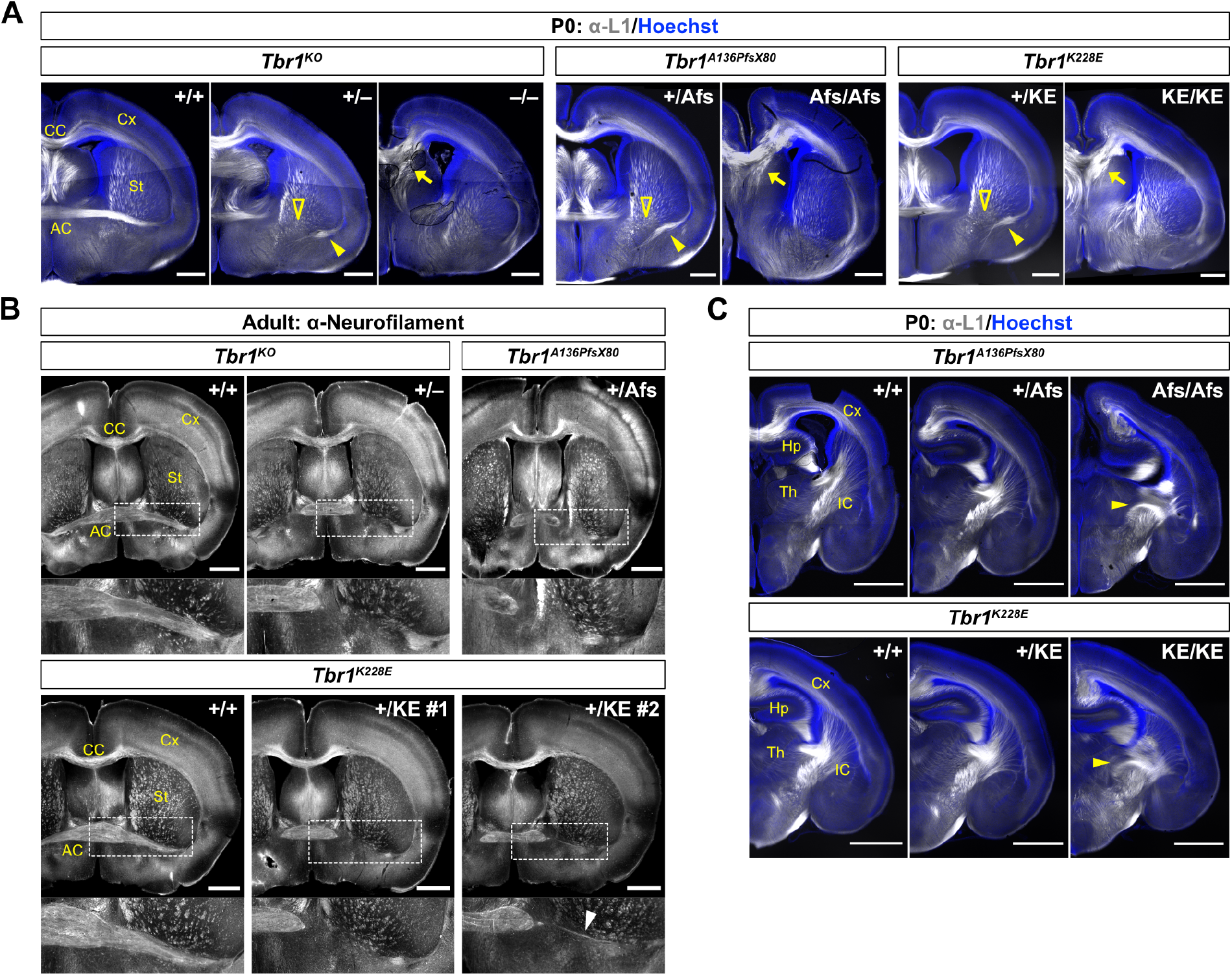
Disruption of Interhemispheric and Subcortical Axon Tracts across *Tbr1* Mutant Mouse Lines. (A) Immunostaining for axon marker L1 (white) and Hoechst nuclear stain (blue) in postnatal day (P) 0 coronal brain sections from *Tbr1* knock-out and patient lines. Open arrowheads indicate absence of posterior limb of anterior commissure (AC) in heterozygotes. Closed arrowheads indicate abnormal external capsule axons in heterozygotes. Arrows indicate bundled and misdirected callosal axons in homozygotes. N = 3-4 mice examined per genotype per line. (B) Immunostaining for axon marker neurofilament (NF) in adult coronal brain sections from *Tbr1* knock-out and patient mutant lines. Inset (dotted lines) shows higher magnification of posterior limb of AC. Arrowhead indicates thin AC posterior limb observed in one *Tbr1^K228E^* heterozygote. N = 3-6 mice examined per genotype per line. (C) Immunostaining for L1 (white) and Hoechst (blue) in P0 coronal brain sections from *Tbr1* patient lines. Arrowheads indicate abnormal organization of internal capsule (IC) axons in homozygotes. AC: anterior commissure, CC: corpus callosum, Cx: cortex, Hp: hippocampus, IC: internal capsule, St: striatum, Th: thalamus. Scale bar = 500 μm in (A); 1 mm in (B) and (C).

In addition to these defects, we identified L1-labeled ectopic axons in the mid-cortex of *Tbr1^−/−^*, *Tbr1^Afs/Afs^*, and *Tbr1^KE/KE^* at P0 (Figure 5A). These axons were constrained to the abnormal inner cortical CUX1+ layer in *Tbr1^−/−^* mice (Figure 5B). Because *Tbr1^−/−^* mice lack the subplate layer along which thalamic axons normally travel to innervate the cortex (Hevner et al., 2001), we sought to determine if these ectopic cortical axons were misguided thalamocortical afferents, or whether they originated intra-cortically. We placed DiI crystals in either the thalamus or the cortex of P0 brains, and we observed DiI-labeled cortical axons forming a narrow mid-cortical tract similar to the L1-labeled ectopic axons (Figure 5B-C). Thalamic axons, on the other hand, were misrouted ventrally into the external capsule and did not enter the cortex (Figure 5D). Altogether, our results show shared axon defects across *Tbr1^KO^* and patient mutant mouse lines, including a newly observed intra-cortical axon defect in homozygous mutants.

**Figure 5.**
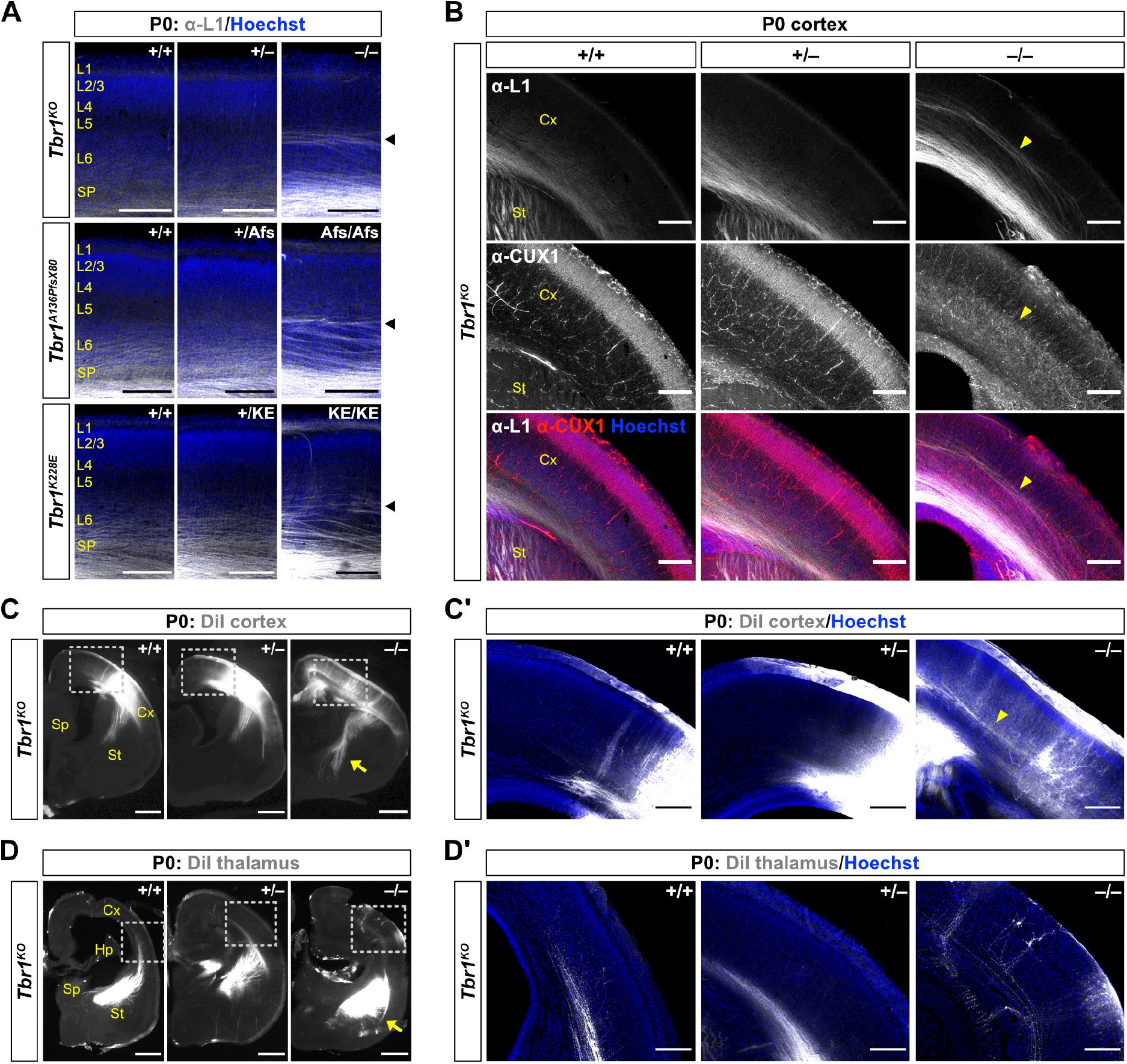
Ectopic Cortical Axons in *Tbr1* Homozygous Mutants Originate within the Cortex. (A) Immunostaining for axon marker L1 (white) and Hoechst nuclear stain (blue) in postnatal day (P) 0 *Tbr1* knock-out and patient line somatosensory cortex. Arrowhead indicates ectopic intracortical axons in homozygotes. (B) Immunostaining for L1, CUX1, and Hoechst in P0 somatosensory cortex from *Tbr1* knock-out line. Yellow arrowheads indicate positioning of ectopic intracortical axons at CUX1+ layer boundary in homozygous mutants. (C-C’) DiI-labeled cortical axons (white) in P0 coronal brain sections from *Tbr1* knock-out line. Arrow indicates subcortical axon overgrowth in homozygous mutants. Dotted lines show inset of cortex in (C’) with Hoechst counterstain (blue) and arrowhead indicating DiI-labeled cortical axons in homozygous mutant. (D-D’) DiI labeled thalamic axons (white) in P0 coronal brain sections from *Tbr1* knock-out line. Arrow indicates misrouted thalamic axons in external capsule. Dotted lines show inset of cortex in (D’) with Hoechst counterstain (blue). Cx: cortex, Hp: hippocampus, Sp: septum, St: striatum, Th: thalamus. Scale bar = 200 μm in (A), (B), (C’), and (D’); 500 μm in (C) and (D).

### *Tbr1* K228E allele is sufficient to prevent apoptosis seen in *Tbr1*-null cortex

Previous investigation of cell survival in *Tbr1^−/−^* cortex found a substantial increase in the apoptotic marker cleaved caspase-3 (CC3) starting by embryonic day (E) 16.5 and continuing to P0 (Bedogni et al., 2010). To compare cell survival upon *Tbr1* deletion versus T-box point mutation, we immunostained for CC3 in *Tbr1^KO^* and *Tbr1^K228E^* mice at P0. In both lines, we saw very few CC3+ cells in WT or heterozygous mutants (Figure 6A-B). Interestingly, we saw drastically increased CC3+ cell density in *Tbr1^−/−^* cortex but not in *Tbr1^KE/KE^* cortex. We posited that the increased apoptosis in *Tbr1^−/−^* could result from misspecification of L6 neurons, which instead acquire L5-like identity with high CTIP2 expression (McKenna et al., 2011). However, when we co-stained for CC3 and CTIP2, we found that CC3+ apoptotic cells were not strictly confined to the CTIP2+ layer (Figure 6C). Furthermore, while CC3+ cells within the CTIP2+ layer showed a neuron-like bipolar morphology, only 16.8% of the total CC3+ cells were CTIP2+ (Figure 6D-E). Thus, complete *Tbr1* knockout causes increased apoptosis in the developing cortex, mostly among non-L5-fated cells, while the *Tbr1* K228E mutation has no impact on cortical cell survival.

**Figure 6.**
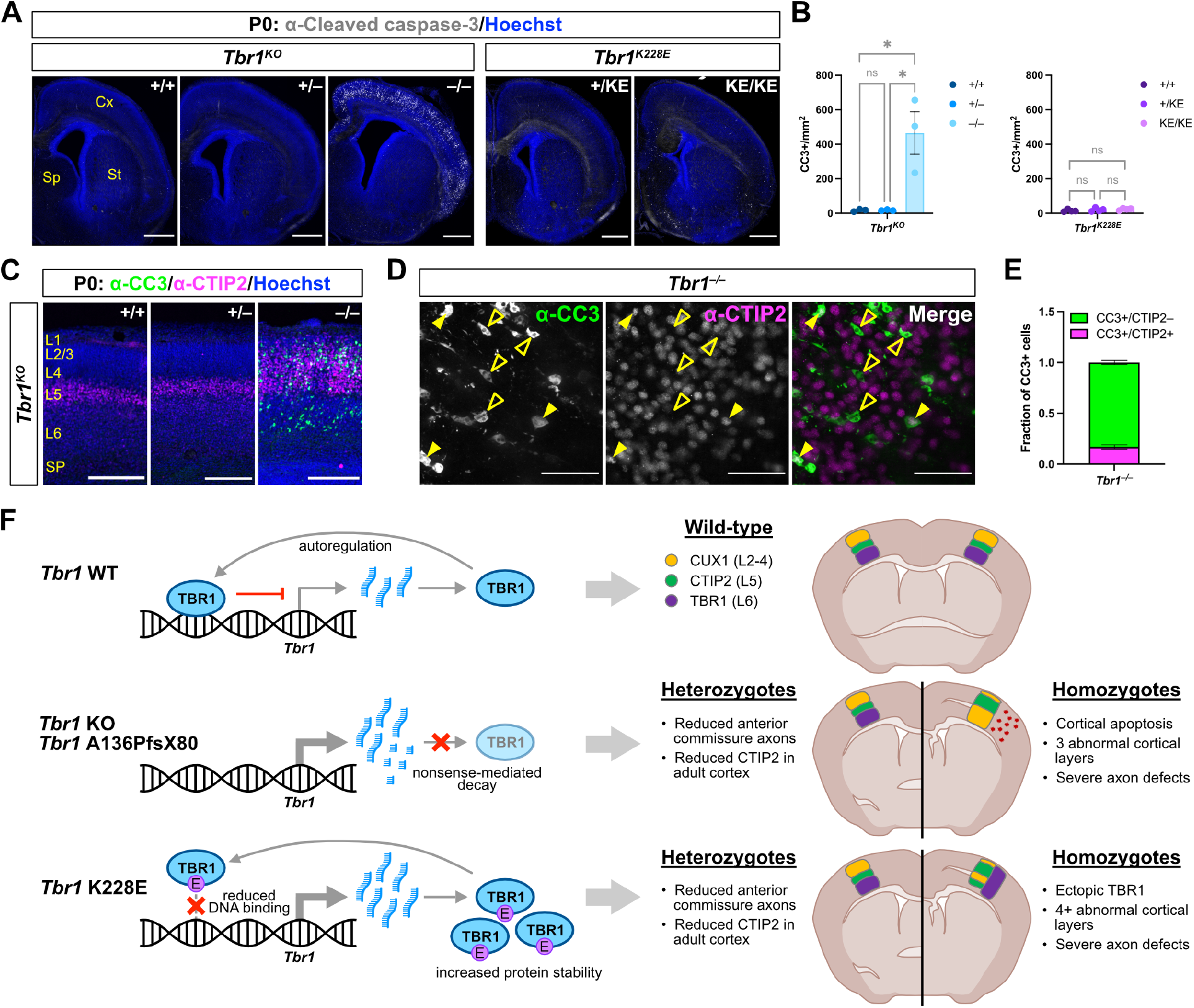
*Tbr1* K228E Allele is Sufficient to Prevent Cortical Apoptosis Seen in *Tbr1* Knock-out Mice. (A-B) Immunostaining and quantification of apoptotic cells expressing cleaved caspase-3 (CC3) (white) in postnatal day (P) 0 cortex from *Tbr1* knock-out and K228E lines (n = 3 mice per genotype). Hoechst nuclear stain is shown in blue. (C) Immunostaining of CC3+ cells (green) in relation to CTIP2+ layers (magenta) in P0 somatosensory cortex of *Tbr1* knock-out line. Hoechst is shown in blue. (D-E) Immunostaining and quantification of CC3+ apoptotic cells (green) co-expressing CTIP2 (magenta) in cortex of P0 *Tbr1^−/−^* mice (n = 3). Open arrowheads indicate CC3+CTIP2– cells; closed arrowheads indicate CC3+CTIP2+ cells. (F) Summary of findings and proposed functional effects of *Tbr1* patient mutations A136PfsX80 and K228E. Scale bar = 500 μm in (A); 200 μm in (C); 50 μm in (D). Data are plotted as mean ± SEM. Each dot represents one animal. One-way ANOVA with Tukey’s multiple comparisons test (B). *p < 0.05; ns, not significant.

## DISCUSSION

In this study, we sought to determine the *in vivo* functional effects of NDD patient-specific *Tbr1* mutations A136PfsX80 and K228E, as well as the in-frame deletion Δ348–353. We saw opposite effects of the patient mutations on TBR1 protein levels, but identified shared phenotypes in heterozygous mutants, including reduction of the anterior commissure and cortical CTIP2 downregulation. We also identified distinct phenotypic effects of homozygous mutations: complete loss of TBR1 caused inversion of the CUX1+ and CTIP2+ cortical layers and extensive apoptosis, while K228E caused cortex-wide TBR1 expression and a complex layering phenotype with no effect on apoptosis (Figure 6F). Unlike these mutations, Δ348–353 did not affect TBR1 levels or cortical layer formation, revealing a portion of the T-box dispensable for these functions. This is the first parallel characterization of multiple patient-specific mutations of *Tbr1 in vivo*, and our results provide several new insights into *Tbr1* function in brain development and disease.

### Construct and face validity of *Tbr1* mouse models for NDDs

Our newly generated *Tbr1^A136PfsX80^* and *Tbr1^K228E^* mouse lines demonstrate a high degree of construct validity for modeling human disorder. With CRISPR/Cas9 genome editing, we generated the homologous patient mutations in the mouse genome without incorporation of artifacts that could impact gene regulation and/or function, such as neomycin cassettes or loxP sites. Furthermore, we included heterozygous mutant mice in our analyses to mimic the heterozygosity of *TBR1* patients (Nambot et al., 2020). We identified A136PfsX80 as a loss-of-function allele based on absence of truncated protein and selectively reduced mutant transcript (Figure 2, S3D). Frameshift mutations producing premature termination codons typically trigger nonsense-mediated decay (NMD) of the mutant transcript in cells, limiting protein production and likely leading to the TBR1 reduction seen in *Tbr1^+/Afs^* mice. We posit that similar early-truncating *TBR1* mutations could also trigger NMD, but *in vivo* validation must be conducted as some transcripts containing premature termination codons have been demonstrated to escape this mRNA surveillance pathway (Holbrook et al., 2004). In contrast with A136PfsX80, we observed a 1.5-fold to 2-fold increase of TBR1 protein in *Tbr1^+/KE^* mice (Figure 2). This mutant protein exhibited impaired DNA binding, increased stability, and altered protein interactions *in vitro* (den Hoed et al., 2018; Deriziotis et al., 2014; Yook et al., 2019). The combination of decreased functionality and retention of certain protein interactions, including homodimerization, opens the possibility of TBR1-K228E acting as a dominant-negative.

Despite functional differences between A136PfsX80 and K228E, we identified a highly penetrant shared phenotype of anterior commissure reduction in heterozygous mutant mice (Figure 4). Thinning or absence of the anterior commissure was also recently observed in a *TBR1* patient cohort, where 7/7 individuals examined by brain MRI demonstrated this phenotype (Nambot et al., 2020). Presence of this defect upon K228E mutation, in which TBR1 retains a degree of activity sufficient for cell survival (Figure 6), further reinforces that anterior commissure development is highly sensitive to mutation of one *Tbr1* copy. This phenotype may prove to be a core biomarker for *TBR1*-related disorders, and its presence in our patient mutant mouse lines demonstrates their face validity as models for these disorders. Along with their construct validity, this could increase their likelihood of high predictive validity in future studies utilizing these models for development and validation of NDD therapies.

While cortical malformations have been reported in some human *TBR1* patients, heterozygous *Tbr1* mutant mice in this and other studies showed grossly normal cortical layer formation (Figure 3) (Huang et al., 2014; Nambot et al., 2020; Vegas et al., 2018; Yook et al., 2019). One possible explanation is that mice are less sensitive to heterozygous *Tbr1* disruption than humans, similar to what was observed for the cortical gene *Dcx* (Nosten-Bertrand et al., 2008). This could result from human-mouse differences in cortical structure (as mice are naturally lissencephalic and may not exhibit *Tbr1*-dependent gyral malformations) or in the magnitude of TBR1 reduction upon gene mutation. Alternatively, cortical malformations in humans may be caused by only a subset of *TBR1* mutations. Supporting this, a close inspection of the human *TBR1* genetic data suggests a correlation between last exon frameshift mutations and cortical gyral abnormalities in MRI (Nambot et al., 2020; Vegas et al., 2018). Independent of cortical formation, neuronal function may be impacted by heterozygous *Tbr1* mutation, as *Tbr1^+/–^* mice show reduced synapses on dendrites of deep-layer glutamatergic neurons (Fazel Darbandi et al., 2020). Whether this could contribute to the abnormal EEG and seizures observed in some *TBR1* patients remains to be determined (Nambot et al., 2020).

Deletion of mouse TBR1 residues 348–353 did not alter TBR1 levels or cortical layer formation (Figure S1). This finding, combined with the low conservation of these residues among mouse T-box proteins, suggests that substitutions or indels within this region may not substantially impact TBR1 function. Thus, the three reported human missense variants T350A, S351R, and P353A may be of low clinical impact as predicted by PolyPhen and SIFT analyses (Karczewski et al., 2020; Landrum et al., 2018). In contrast, the two human nonsense mutations identified in this region show higher pathogenicity predictions, and the TBR1-S351X protein was verified to be dysfunctional *in vitro* (Deriziotis et al., 2014). While TBR1-Δ348–353 functionality is sufficient for corticogenesis, we cannot rule out subtler effects of this deletion on cortical development or neuronal function.

### Negative autoregulation of *Tbr1* expression in early postnatal cortex

Our analyses of *Tbr1* expression in multiple mutant mouse lines provide insights into regulation of this gene in early postnatal cortex. In *Tbr1^KO^* and *Tbr1^A136PfsX80^* mutants, we observed a mismatch of high *Tbr1* transcript levels but lowered TBR1 protein levels, while in *Tbr1^K228E^* mutants, we observed increases of both transcript and protein (Figure 2). Considering that transcription factors commonly perform autoregulatory functions to ensure proper abundance in cells (Crews and Pearson, 2009), we propose a mechanism of negative autoregulation by TBR1 in postnatal cortex (Figure 6F). Previous chromatin immunoprecipitation with sequencing (ChIP-seq) assays in developing mouse cortex identified several TBR1-bound genomic sites near the *Tbr1* gene, which could act as potential autoregulatory elements (Fazel Darbandi et al., 2018; Notwell et al., 2016). Following this model, in *Tbr1^KO^* and *Tbr1^A136PfsX80^* cortex, mutant transcripts would undergo NMD leading to TBR1 protein reduction, which would in turn derepress *Tbr1* transcription. In *Tbr1^K228E^* cortex, the TBR1-K228E protein would exhibit reduced DNA binding (Yook et al., 2019), also leading to failure of autorepression. However, these transcripts would not undergo NMD and instead lead to overabundance of protein, which could be further exacerbated by the increased stability of the mutant protein observed *in vitro* (den Hoed et al., 2018; Yook et al., 2019).

Failure of TBR1 autorepression might also contribute to the unexpected ectopic TBR1 in nearly all cortical neurons of *Tbr1^KE/KE^* mice (Figure 2, S3). It has been postulated that the transcription factor sequence PAX6 → EOMES (TBR2) → TBR1 is required for differentiation of cortical radial glia into intermediate progenitors and then postmitotic neurons (Englund et al., 2005). This was recently supported by temporal single-cell RNA-seq showing that progenitors born at E12 (eventual deep-layer) through E15 (eventual upper-layer) express *Tbr1* upon neuronal differentiation (http://genebrowser.unige.ch/telagirdon/) (Telley et al., 2019). Following this obligatory *Tbr1* activation, newborn neurons must either maintain or repress its expression depending on their subtype specification. For instance, TBR1 is maintained in L6 neurons postnatally, while L5 corticospinal fate requires direct repression of *Tbr1* by CTIP1 (BCL11A) (Canovas et al., 2015). If TBR1 autorepression also contributes to this developmental layer patterning, its failure in *Tbr1^KE/KE^* mice could then lead to high pan-neuronal *Tbr1* expression. Exploring this phenotype could lend important insights into the temporal regulation of transcription factor expression and neuronal fate specification in the developing cortex. Additionally, enhanced understanding of *Tbr1* regulation, including autoregulation, could be an important consideration for therapeutic avenues involving correction of WT TBR1 protein levels in patients with *TBR1* mutations.

### Insights into the multifaceted roles of *Tbr1* in corticogenesis

Previous studies have demonstrated that *Tbr1* mediates proper organization of cortical neurons: *Tbr1^−/−^* mice show *reeler*-like cortical layering and abnormal distribution of interneurons, while *Tbr1^+/KE^* mice exhibit subtle displacement of parvalbumin-expressing interneurons in the medial prefrontal cortex (mPFC) (Hevner et al., 2001; Huang et al., 2014; Yook et al., 2019). As *Tbr1* is not normally expressed in interneurons, defects in this cell type are considered secondary to defects in *Tbr1*-expressing cell types (Hevner et al., 2001). With our assessment of layer marker fluorescence profiles, we found that homozygous *Tbr1* mutants formed abnormal layers in distinct manners depending on the mutation (Figure 3). This could result from differential impacts of the mutations on Cajal-Retzius cells, which express *Tbr1* and secrete Reelin in early cortical development to establish proper lamination (Hevner et al., 2001). Future studies could disentangle the contributions of projection neuron *Tbr1* versus Cajal-Retzius *Tbr1* to cortical layer formation using conditional knockout approaches. In contrast to homozygotes, we quantitatively confirmed nearly identical layer positions between WT and heterozygotes in each line (Figure 3). We also did not observe changes in the distribution of parvalbumin-or somatostatin-expressing interneurons in heterozygotes (Figure S4). While our result differs from the prior interneuron finding in *Tbr1^+/KE^* mice, it could be explained by differences in cortical area examined, as we performed measurements in primary somatosensory cortex rather than mPFC (Yook et al., 2019). Considering *Tbr1*’s function in establishing frontal areal identity (Bedogni et al., 2010), its indirect roles in cortical interneuron migration could be more pronounced in the mPFC.

*Tbr1* is also critical for the specification and maintenance of L6 corticothalamic neuronal fate. *Tbr1^−/−^* mice show conversion of L6-fated neurons to abnormal L5-like corticospinal neurons, while L6-specific deletion of *Tbr1* after L6 specification causes upregulation of L5 marker genes and incomplete corticothalamic axonal arborization (Fazel Darbandi et al., 2020; Fazel Darbandi et al., 2018; Han et al., 2011; McKenna et al., 2011). These and other studies have elucidated the following mechanisms for deep-layer neuronal fate specification: L6 corticothalamic fate is established by TBR1 directly repressing *Fezf2*, while L5 corticospinal fate is established by CTIP1 (BCL11A) directly repressing *Tbr1*, and FEZF2 directly activating *Ctip2* (*Bcl11b*) (Canovas et al., 2015; Chen et al., 2005a; Chen et al., 2008; Chen et al., 2005b; Molyneaux et al., 2005). In our *Tbr1* mutant mice, we saw a seemingly paradoxical effect on CTIP2 levels, which were increased in homozygous mutants but decreased in adult heterozygous mutants (Figure 3). One potential explanation is weak but direct activation of *Ctip2* by TBR1 in a cell-autonomous manner. ChIP-seq in E15.5 cortex identified five TBR1-bound sites in the vicinity of *Ctip2*, and *Tbr1^−/−^* mice show decreased CTIP2 levels in E14.5 cortical plate (Bedogni et al., 2010; Notwell et al., 2016). Furthermore, in *Fezf2^−/−^* mice, CTIP2 is absent from L5 but maintained in L6, where it colocalizes with TBR1 (Molyneaux et al., 2005). Direct *Ctip2* activation by TBR1 could explain the CTIP2 downregulation seen in our adult heterozygous *Tbr1* mutant mice. Conversely, in homozygous mutants, the derepression of *Fezf2* over several days of neurogenesis would supersede the direct effects of TBR1 loss on *Ctip2* expression, leading to overabundance of CTIP2+ neurons by P0. These results further highlight the complex and nuanced regulatory networks among transcription factors in establishing cortical projection neuron identity.

Lastly, *Tbr1* is essential for proper cortical axon development. In *Tbr1^−/−^* mice, early-born neurons send their axons to the spinal cord rather than to the thalamus as a consequence of the L6→L5 neuronal fate-switch (Han et al., 2011; McKenna et al., 2011). These mice also show corpus callosum defects, with some axons crossing the midline but others forming errant disorganized bundles described as Probst bundles (Hevner et al., 2001). We identified an additional defect in homozygous *Tbr1* mutant mice where cortical axons form a distinct tract traveling along the upper boundary of the abnormal CUX1+ layer (Figure 5). The mechanism(s) for these intracortical axon defects remains to be determined, but one possible explanation is that TBR1 acts cell-autonomously within intracortical projection neurons to ensure proper axon targeting. Both retrograde tracing and genetic labeling have shown that *Tbr1* is expressed in L6 intracortical neurons (Galazo et al., 2016; Matho et al., 2021; Tasic et al., 2016). Alternatively, these axon defects could be secondary to the layering defects in these mice, as extracellular components important for axon guidance could be abnormally distributed. In addition to anterior commissure defects, a subset of human *TBR1* patients show callosal or other white matter defects in MRI, underscoring the need to understand TBR1-dependent axon guidance mechanisms in the context of disease. More broadly, our identification of shared and distinct cortical phenotypes across *Tbr1* mouse models highlights the utility of *in vivo* characterization of patient-specific mutations to elucidate genotype-phenotype relationships in NDDs.

## AUTHOR CONTRIBUTIONS

This project was conceived by M.C., R.A.B., A.C.A., K.M.W., and B.J.O. Experimental design was performed by M.C. and R.A.B. New mouse lines were generated by L.M.F. Data collection and analyses were performed by M.C., J.N.J., and S.G. Biological interpretation was performed by M.C., A.C.A., K.M.W., and B.J.O. The manuscript was written by M.C. with input from all authors.

## ACKNOWLEDGMENTS

We thank current and former members of the O’Roak, Wright, and Adey lab groups for their comments and technical support for this study, especially Dominica Cao, Brooke DeRosa, Sara Evans-Dutson, Bridget Fitzgerald, Destine Krenik, Cierra LeBlanc, Neville Lee, Amanda Mar, Ryan Mulqueen, Andrew Nishida, and Lindsay Wourms. We also thank current and past members of the BRAINS award advisory committee for their advice and mentorship: Eric Fombonne, Marc Freeman, Gail Mandel, Tomasz Nowakowski, and Soo-Kyung Lee. We also thank Yingming Wang at the OHSU Transgenic Mouse Models Core for technical assistance with generating mouse lines, as well as initial support from an OHSU University Shared Resources Pilot Award (B.J.O). Research reported in this publication was supported by the National Institute of Mental Health of the National Institutes of Health under award number R01MH113926 (B.J.O). The content is solely the responsibility of the authors and does not necessarily represent the official views of the National Institutes of Health.

## DECLARATION OF INTERESTS

The authors declare no competing financial interests.

## METHODS

### RESOURCE AVAILABILITY

#### Lead contact

Further information and requests for resources and reagents should be directed to and will be fulfilled by the lead contact, Brian J. O’Roak.

#### Materials availability

Mouse lines generated in this study will be made available through the Jackson Laboratory.

#### Data and code availability

All data reported in this paper, and any additional information required to reanalyze the data, is available from the lead contact upon request. This paper does not report original code.

### EXPERIMENTAL MODEL AND SUBJECT DETAILS

#### Animals

*Tbr1^KO^* mice (Bulfone et al., 1998) were rederived from cryopreserved sperm obtained from MMRRC at UC Davis (030263-UCD) and backcrossed to a C57BL/6 background for at least two generations prior to data collection. *Tbr1^A136PfsX80^*, *Tbr1^K228E^*, and *Tbr1^Δ348–353^* mice were generated on a C57BL/6NJ background and backcrossed to C57BL/6NJ for at least two generations prior to data collection. Mice of both sexes were used for all experiments. For molecular and histological experiments, mice were analyzed at P0 or adulthood (9-39 weeks). For weight and behavior assessments, mice were analyzed at P4, P7, P10, and P14. Cohorts for each experiment were comprised of littermate wild-type (WT) and mutant mice from at least 2 separate litters. Mice were group housed under a 12 h light/dark cycle and given *ad libitum* access to food and water. All animal procedures were approved by Oregon Health & Science University Institutional Animal Care and Use Committee.

### METHOD DETAILS

#### Generation of mouse lines

CRISPR sgRNA design to generate *Tbr1^A136PfsX80^*, *Tbr1^K228E^*, and *Tbr1^Δ348–353^* founders was performed using the CRISPOR tool (http://crispor.tefor.net/) (Concordet and Haeussler, 2018). Generation of mutant mice was performed by the Oregon Health & Science University Transgenic Mouse Models Core based on (Aida et al., 2015). C57BL/6NJ zygotes were co-injected with Cas9 protein (50 ng/μl) (New England Biolabs) or *Cas9* mRNA (1000 ng/μl) (TriLink BioTechnologies), sgRNA (30 ng/μl) (Synthego), and DNA donor (100 ng/μl) (Integrated DNA Technologies) containing the patient mutation. See Table S1 for sgRNA and donor sequences. Embryos were transplanted into pseudopregnant recipient female CD-1 mice, and founders from these litters were identified via Sanger sequencing.

#### Mouse genotyping

*Tbr1^KO^* mice were PCR genotyped using primers amplifying genomic *Tbr1* and the neomycin cassette. *Tbr1^A136PfsX80^* and *Tbr1^K228E^* mice were genotyped using PCR amplification of the mutation-containing genomic region followed by restriction enzyme digest of the PCR product (BtsCI for A136PfsX80, BtsIMutI for K228E). *Tbr1^Δ348–353^* mice were PCR genotyped using primers amplifying genomic *Tbr1*. See Table S1 for primer sequences.

#### Neonatal weight and motor assessments

Weight measurements and negative geotaxis testing were performed at P4, P7, P10, and P14 as previously described (Hill et al., 2008). For negative geotaxis, each pup was placed with its head pointing downward on a 45° incline, and the latency for the pup to face upward on the incline was recorded. If the pup failed to turn within 60 seconds, or if the pup fell down the incline 3 times, the latency was recorded as 60 seconds.

#### Western blot

Cortex was dissected at P0 or adulthood (10-39 weeks), flash frozen, and stored at −80°C until all samples were collected. Frozen tissue from one cortical hemisphere per mouse (~30 mg at P0, ~100 mg at adulthood) was dounce homogenized in ice-cold RIPA buffer (150 mM NaCl, 1% IGEPAL CA-630, 0.5% sodium deoxycholate, 0.1% SDS, 50 mM Tris-HCl pH 8.0, 2 mM EDTA pH 8.0) containing Protease Inhibitor Cocktail (Roche). Nuclei were lysed using a Misonix XL-2000 Probe Sonicator with 5-10 second ON / 20 second OFF intervals until the lysate was clear (2 rounds for P0, 4-5 rounds for adult). Lysates were further incubated on ice for 30 minutes then centrifuged at 10,000 *rcf* for 10 minutes at 4°C to remove debris. Protein concentrations were determined using the Pierce BCA Protein Assay Kit (ThermoFisher). Laemmli SDS sample buffer (Alfa Aesar) was added to lysates which were boiled at 95°C for 5 minutes prior to SDS-PAGE. Total protein (20-30 μg per sample) was resolved on 4-15% polyacrylamide gels (Bio-Rad) and transferred to PVDF membranes. Membranes were incubated in block solution (5% milk in TBS with 0.1% Tween-20 [TBST]) for 1 hour at room temperature, incubated in primary antibodies in block solution overnight at 4°C, washed in TBST 4X 10 minutes, incubated in secondary antibodies in block solution for 1 hour at room temperature, washed in TBST 4X 10 minutes, and imaged with an Odyssey CLx using Image Studio software (LI-COR Biosciences). Antibodies and dilutions used are listed in the Key Resources Table.

#### RT-qPCR and Sanger sequencing

Cortex was dissected at P0, flash frozen, and stored at −80°C until all samples were collected. Frozen tissue from one cortical hemisphere (~30 g) was lysed in 1 ml TRIzol Reagent (Invitrogen) and homogenized by passing through a 25G needle. Total RNA was extracted using the RNeasy Lipid Tissue Mini Kit (QIAGEN) with on-column DNase digestion with RNase-Free DNase Set (QIAGEN) according to the manufacturer’s protocols. cDNA was synthesized from 1 μg total RNA using the ProtoScript® II First Strand cDNA Synthesis Kit (NEB) with oligo-dT priming according to the manufacturer’s protocol. For RT-qPCR experiments, cDNA templates and no RT controls were diluted 1:20 for multiplexed PrimeTime qPCR Assays using predesigned *Tbr1* and *Actb* primers with FAM and SUN probes, respectively (Integrated DNA Technologies). Probe fluorescence was measured with a CFX Connect Real-Time PCR Detection System using CFX Manager Software (Bio-Rad) running the following cycling program: 95°C for 3 minutes, 40 cycles of 95°C 15 seconds and 60°C 1 minute, 4°C hold. RT-qPCR primer efficiencies for *Tbr1* and *Actb* were calculated as 98.5% and 100.4% respectively using WT cDNA for the standard curve (undiluted to 1:10,000). For RT-PCR and Sanger sequencing in *Tbr1^A136PfsX80^* mice, 1 μl cDNA was used as PCR template for M13 sequence-containing primers spanning exons 1 and 2, and purified PCR product was Sanger sequenced using M13-forward primer. See Table S1 for primer sequences.

#### Immunohistochemistry

For P0 samples, whole brains were drop-fixed in 4% EM-grade paraformaldehyde (PFA) overnight at 4°C, then washed in PBS. For adult samples (9-19 weeks), brains were fixed via transcardial perfusion with 10 ml PBS and 10 ml 4% PFA, followed by post-fix in 4% PFA for 1 hour at 4°C, followed by PBS wash. For cortical layering, interneuron, axon, and apoptosis marker IHC, brains were embedded in 3% LMP agarose and sectioned coronally at 100 μm using a Leica VT1200 S vibratome. Free-floating sections anterior to the hippocampus were incubated in primary antibodies in block solution (2% normal donkey serum, 0.2% Triton X-100 in PBS) for 48-72 hours at 4°C, washed in PBS for 5+ hours, incubated in secondary antibodies and Hoechst stain 1:5000 in block solution overnight at 4°C, washed in PBS for 5+ hours, mounted onto glass slides, and coverslipped with Fluoromount-G (Southern Biotech). For TBR1/NeuN IHC in *Tbr1^K228E^* cortex, fixed brains were cryoprotected in 30% sucrose overnight at 4°C, washed in PBS, embedded in Tissue-Tek OCT compound, and sectioned coronally at 20 μm using a Tanner TN50 cryostat. Free-floating cryosections were incubated in primary antibodies in block solution overnight at 4°C, washed in PBS 3X 10 minutes, incubated in secondary antibodies and Hoechst stain 1:5000 in block solution for 1 hour at room temperature, washed in PBS 3X 10 minutes, mounted onto glass slides, and coverslipped with Fluoromount-G (Southern Biotech). Antibodies and dilutions used are listed in the Key Resources Table.

#### DiI labeling

P0 whole brains were drop-fixed in 4% EM-grade PFA for at least 24 hours at 4°C, then washed in PBS. For corticothalamic labeling, DiI crystals were embedded in S1 cortex along the anteroposterior axis. For thalamocortical labeling, DiI crystals were embedded in thalamus after removal of hindbrain. Labeled brains were incubated in 4% PFA at 37°C until labeling was visible in the target brain regions or axon tracts (~9 days). Brains were embedded in 3% LMP agarose and sectioned coronally at 150 μm using a Leica VT1200 S vibratome. Sections were stained with Hoechst 1:5000 in PBS for 10 minutes, mounted onto glass slides, and coverslipped with Fluoromount-G (Southern Biotech).

#### Fluorescence image acquisition

Images were acquired using Zeiss ZEN software. IHC sections were imaged on a Zeiss Axio Imager M2 upright microscope equipped with an ApoTome2. For cortical layering, interneuron, and apoptosis IHC, Z-stacks were obtained through the tissue section using the optimal interval for each objective. For axon IHC, images were obtained using Tile Scan mode and stitched using ZEN. DiI-labeled sections were imaged on a Zeiss Axio Zoom.V16 dissecting microscope.

### QUANTIFICATION AND STATISTICAL ANALYSIS

#### Western blot

Western blot band intensities were measured using “Analyze>Gels” in ImageJ2/FIJI software (Rueden et al., 2017). Within each blot, each TBR1 signal was normalized to its corresponding loading control signal, and then each normalized TBR1 signal was adjusted to the average normalized TBR1 signal across WT replicates.

#### RT-qPCR and Sanger sequencing

For RT-qPCR, Ct values were obtained using the “single threshold” Cq Determination Mode in CFX Manager Software (Bio-Rad). *Tbr1* fold gene expression was calculated using the 2^−ΔΔCt^ method with *Actb* as the reference gene and adjustment to the WT average. Sanger sequencing traces were visualized using Sequencher software (Gene Codes Corporation).

#### Cortical layering

Cortical layering analyses were performed on Z-projection images of coronal sections using ImageJ2/FIJI software. Equivalent background subtraction and brightness/contrast adjustments were applied to all images within an experiment. Images were rotated until layers in S1 were roughly horizontal, then a 1024 × 1800 pixel rectangle was drawn over S1 and rescaled to encompass layer 1 through subplate. Pixel intensities were averaged horizontally within the rectangular selection using “Analyze>Plot Profile”, and then these values were averaged into 100 equal bins from layer 1 through subplate (% cortical distance). Binned values for each sample were min-max normalized using the minimum and maximum average bin values across WT replicates. For binning of “% cortical distance” into layers for CTIP2 fluorescence comparisons, the following bins were determined based on Hoechst fluorescence: L6: 0%–35%, L5: 36%–55%, L2-4: 56%–100%.

#### Interneuron distribution

Interneuron distribution analyses were performed on Z-projection images of coronal sections using ImageJ2/FIJI software. Equivalent background subtraction and binary thresholds were applied to all images within an experiment. Images were rotated until layers in S1 were roughly horizontal, then a rectangular ROI measuring 1024 pixels wide × (height in pixels of layer 1 through subplate) was drawn over S1. Within this ROI, interneuron counts and X-Y coordinates were obtained using “Measure>Analyze Particles” with equivalent particle size parameters for all images. These values were then used to determine number of cells per ROI, fraction of cells per bin (10 bins or 2 bins), and cumulative density of cells along the ROI.

#### Apoptosis

Apoptosis analyses were performed on Z-projection images of coronal sections using ImageJ2/FIJI software. Cells within one cortical hemisphere of one section per animal were manually counted using “Plugins>Analyze>Cell Counter”. Cortical area was calculated using the Polygon tool to manually select the ROI.

#### Statistics

Data plotting and statistical tests were performed using Prism 9 (GraphPad Software). Data are represented as means ± SEM or as boxplots. Each dot represents one animal where applicable. Analyses between two groups were performed using unpaired t-tests. Analyses between three groups were performed using one-way ANOVA with Tukey’s multiple comparisons or two-way ANOVA with Šidák’s multiple comparisons test. Analysis of postnatal weight and motor assessment was performed using two-way repeated measures ANOVA with Šidák’s multiple comparisons test. Analysis of cumulative interneuron distribution was performed using Kolmogorov-Smirnov tests. Significance was defined as p < 0.05.

## Notes

### Competing Interest Statement

The authors have declared no competing interest.

