## Supplemental Material for "Shared and distinct functional effects of patient-specific *Tbr1* mutations on cortical development"

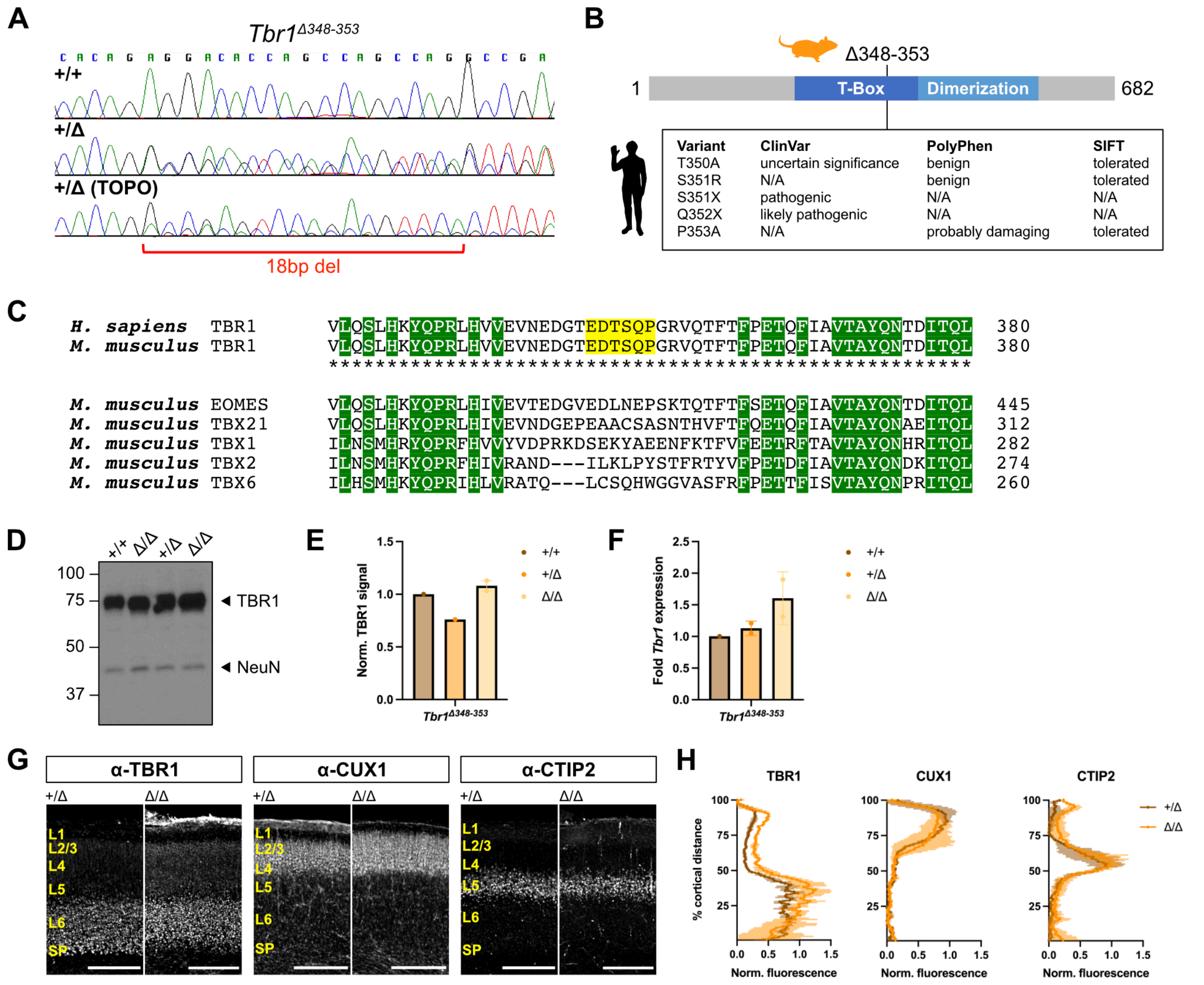


**Supplemental Figure 1. In-Frame Deletion Identifies Dispensable Region in T-Box Domain of TBR1**
(A) Sanger sequencing of wild-type and CRISPR-generated *Tbr1* mutant genomic DNA showing an 18-base-pair deletion, which causes in-frame deletion of amino acids 348–353 (Δ348–353).
(B) Schematic of TBR1 protein showing location of Δ348–353 in T-box domain (top) and human mutations located within this region from ClinVar and gnomAD databases (bottom).
(C) Multiple sequence alignment of residues 348–353 (yellow highlight) and the surrounding region between human and mouse TBR1 (top) and across mouse T-box family proteins (bottom). Asterisks indicate conserved residues between human and mouse TBR1. Green highlight indicates conserved residues across all proteins.
(D-E) Western blot for TBR1 and quantification in cortical lysates from postnatal day (P) 0 *Tbr1*$^{Δ348–353}$ mice (n = 1-2 mice per genotype).
(F) RT-qPCR for *Tbr1* expression levels in P0 *Tbr1*$^{Δ348–353}$ cortex (n = 1-2 mice per genotype).
(G-H) Immunostaining and fluorescence quantification of TBR1, CUX1 (L2-4), and CTIP2 (L5) in P0 *Tbr1*$^{Δ348–353}$ somatosensory cortex (n = 2 mice per genotype).
CC: corpus callosum, Cx: cortex, L: layer, SP: subplate, Th: thalamus. Scale bar = 200 µm in (G); 500 µm in (I) and (J). Data are plotted as mean ± SEM. Each dot represents one animal.

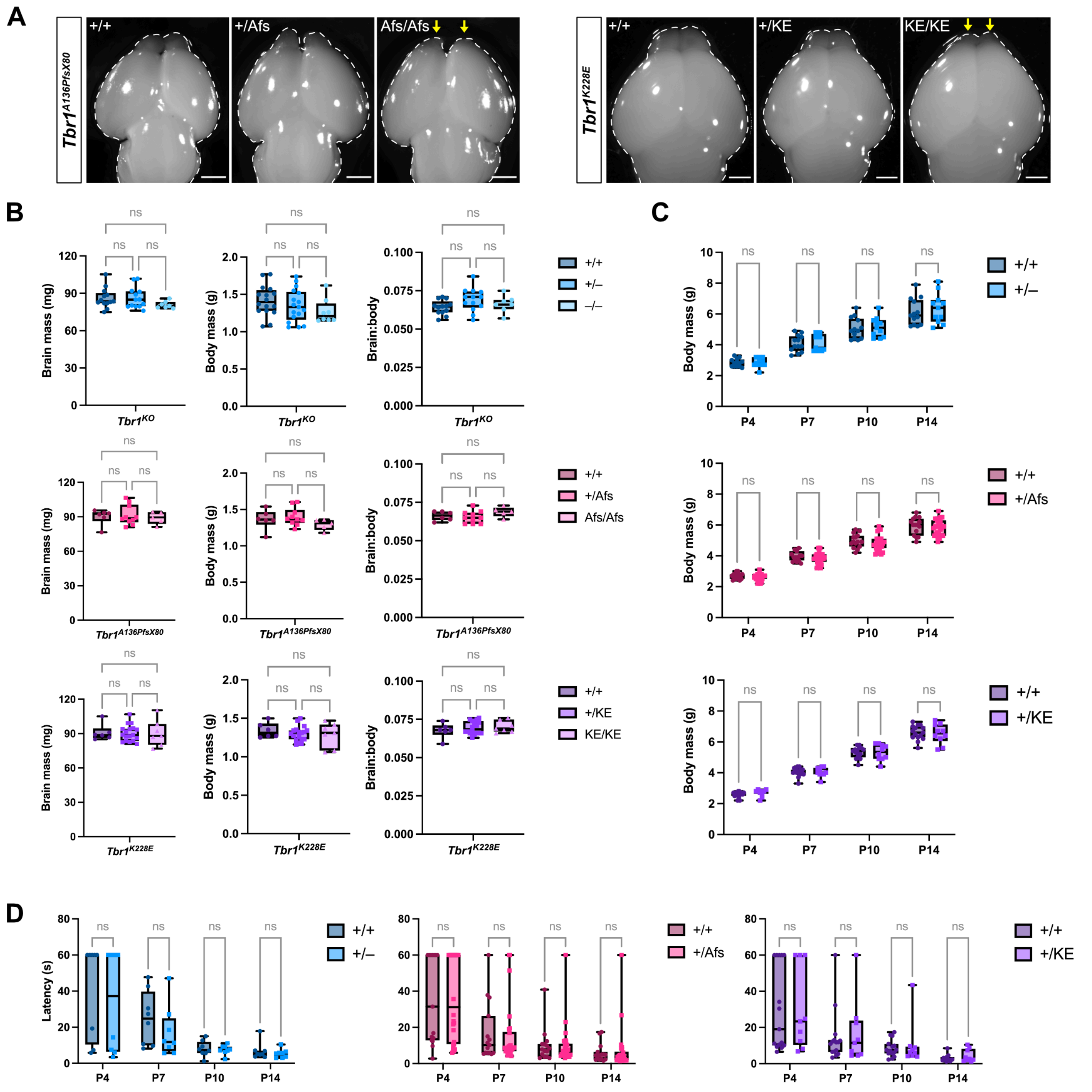


**Supplemental Figure 2. Assessment of Postnatal Brain and Body Development in *Tbr1* Mutant Mouse Lines**

(A) Dorsal view of brains from embryonic day 15.5 *Tbr1*$^{A136PfsX80}$ mice and postnatal day (P) 0 *Tbr1*$^{K228E}$ mice. Yellow arrows indicate underdeveloped olfactory bulbs in homozygous mutants.

(B) Brain mass, body mass, and brain-to-body mass ratio measurements for *Tbr1* knock-out and patient mutant mouse lines at P0 (n = 5-18 mice per genotype).

(C) Body mass measurements for *Tbr1* knock-out and patient mutant mouse lines across postnatal development (n = 10-27 mice per genotype).

(D) Latency for mouse pups to orient upward on a slope in the negative geotaxis test of motor coordination (n = 8-26 mice per genotype).

Scale bar = 1 mm in (A). Data are plotted as mean ± SEM. Each dot represents one animal. One-way ANOVA with Tukey's multiple comparisons test (B); two-way repeated measures ANOVA with Šídák's multiple comparisons test (C) and (D). ns, not significant.

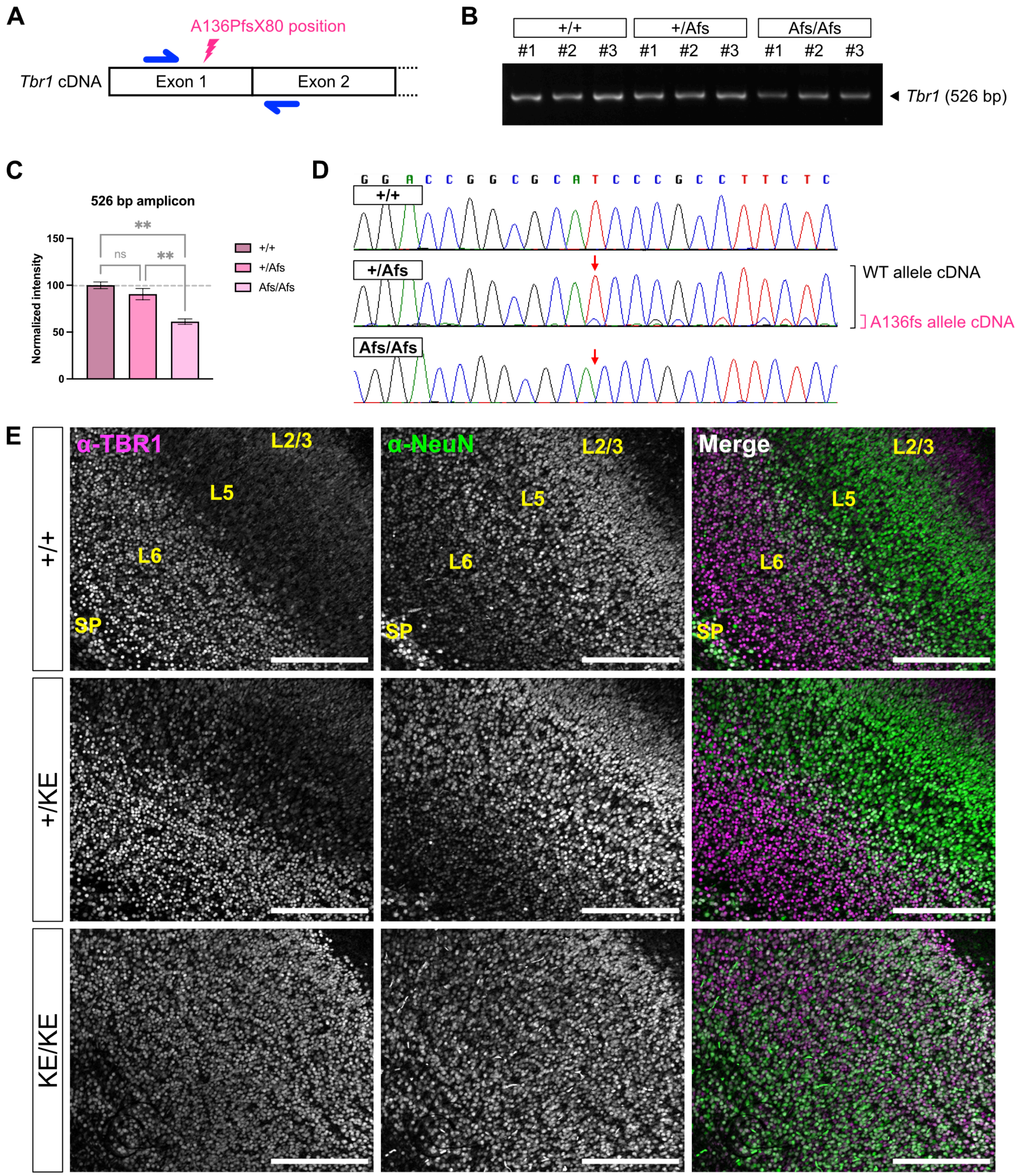


**Supplemental Figure 3. Additional Analysis of *Tbr1* Expression in Patient Mutant Mouse Lines**
(A) Schematic of *Tbr1* cDNA exons 1 and 2 showing positions of A136PfsX80 mutation (magenta) and PCR primers (blue).
(B-C) RT-PCR for *Tbr1* using primers from (A) in postnatal day (P) 0 $Tbr1^{A136PfsX80}$ cortex (n = 3 mice per genotype).
(D) Sanger sequencing of *Tbr1* amplicons generated in (B). Red arrows indicate position of the thymine (T) nucleotide deletion producing p.A136PfsX80. Brackets indicate peaks corresponding to WT (black) and mutant (magenta) allele cDNA in heterozygote.
(E) Immunostaining for TBR1 and NeuN in P0 $Tbr1^{K228E}$ somatosensory cortex.
Scale bar = 200 µm in (E). Data are plotted as mean ± SEM. One-way ANOVA with Tukey's multiple comparisons test (C). **p < 0.01; ns, not significant.

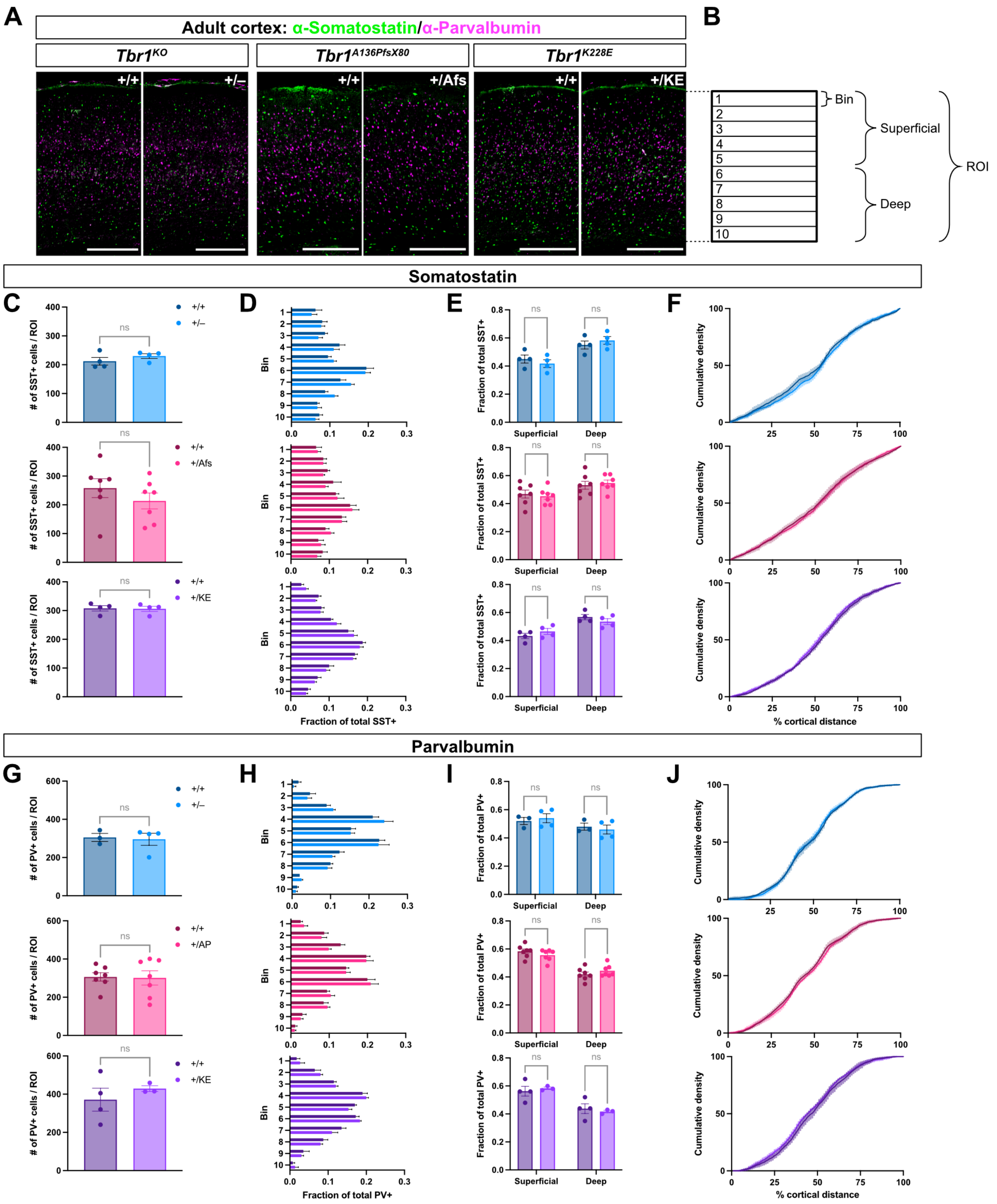


**Supplemental Figure 4. Distribution of Cortical Interneurons in *Tbr1* Patient Mutant Mice**

(A) Immunostaining for interneuron subtype markers somatostatin (SST, green) and parvalbumin (PV, magenta) in adult *Tbr1* knock-out and patient mutant line somatosensory cortex.

(B) Schematic illustrating bins and region-of-interest (ROI) for interneuron quantification.

(C) Number of SST+ cells per ROI (n = 3-7 mice per genotype).

(D) Fraction of SST+ cells distributed across ten equal-sized bins of ROI.

(E) Fraction of SST+ cells distributed between superficial (bins 1-5) and deep (bins 6-10) cortex.

(F) Cumulative density of SST+ cells across the cortical mantle.

(G-J) Quantification of PV+ cells as in (B-E) (n = 3-7 mice per genotype).

Scale bar = 500 μm in (A). Data are plotted as mean ± SEM. Each dot represents one animal. Unpaired Student's t-test (C) and (G); two-way ANOVA with Šídák's multiple comparisons test (D), (E), (H), and (I); Kolmogorov-Smirnov test (F) and (J). ns, not significant.

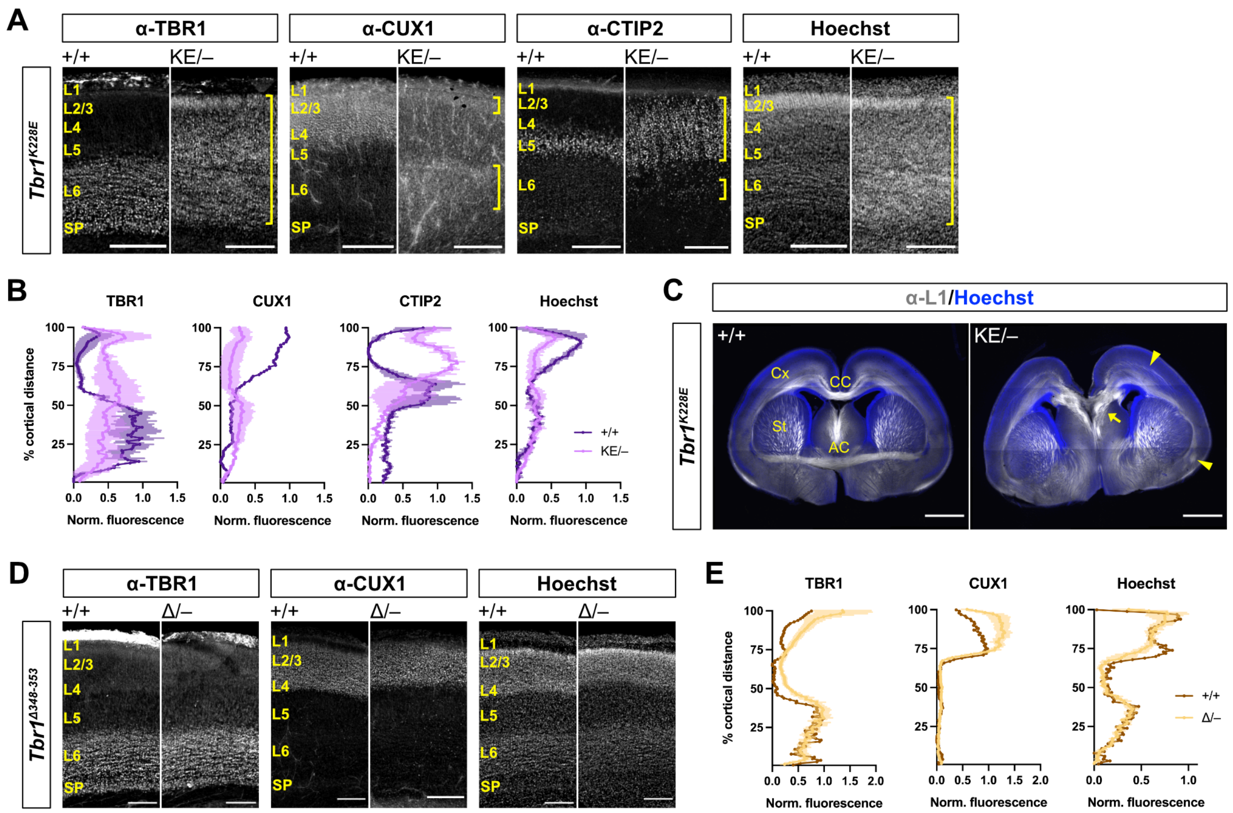


**Supplemental Figure 5. *Tbr1* K228E Fails to Complement *Tbr1* Knock-out Allele**

(A-B) TBR1, CUX1, and CTIP2 immunostaining and Hoechst nuclear stain in somatosensory cortex of P0 offspring from *Tbr1*$^{+/K228E}$ and *Tbr1*$^{+/–}$ complementation crosses (n = 1-2 mice per genotype). Yellow brackets indicate abnormal cortical layers formed in *Tbr1*$^{K228E/–}$ mice.

(C) Immunostaining for axon marker L1 (white) and Hoechst stain (blue) in coronal brain sections of P0 offspring from *Tbr1*$^{+/K228E}$ and *Tbr1*$^{+/–}$ crosses. Arrowheads indicate abnormal cortical axons in *Tbr1*$^{K228E/–}$ mutant. Arrow indicates callosal Probst bundles in *Tbr1*$^{K228E/–}$ mutant.

(D-E) TBR1, CUX1, and Hoechst staining and fluorescence quantification in somatosensory cortex of P3 offspring from *Tbr1*$^{+/Δ348–353}$ and *Tbr1*$^{+/–}$ complementation crosses (n = 1-5 mice per genotype).

AC: anterior commissure, CC: corpus callosum, Cx: cortex, L: layer, SP: subplate, St: striatum. Scale bar = 200 μm in (A) and (D); 1 mm in (C). Data are plotted as mean ± SEM.

**Key Resources Table**

| REAGENT or RESOURCE | SOURCE | IDENTIFIER |
|---|---|---|
| **Antibodies** | | |
| Rabbit anti-β-Tubulin III (WB 1:5000) | Sigma-Aldrich | Cat# T2200, RRID:AB_262133 |
| Rabbit anti-Cleaved Caspase-3 (Asp175) (IHC 1:500) | Cell Signaling Technology | Cat# 9661, RRID:AB_2341188 |
| Rat anti-CTIP2 (25B6) (IHC 1:500) | Abcam | Cat# ab18465, RRID:AB_2064130 |
| Rabbit anti-CDP/CUX1 (M-222) (IHC 1:500) | Santa Cruz Biotechnology | Cat# sc-13024, RRID:AB_2261231 |
| Rat anti-L1 (clone 324) (IHC 1:500) | Millipore | Cat# MAB5272, RRID:AB_2133200 |
| Mouse anti-NeuN (1B7) (IHC 1:500) | Abcam | Cat# ab104224, RRID:AB_10711040 |
| Mouse anti-NeuN (clone A60) (WB 1:200) | Millipore | Cat# MAB377, RRID:AB_2298772 |
| Mouse anti-Neurofilament/NF-M (IHC 1:500) | Developmental Studies Hybridoma Bank | Cat# 2H3, RRID:AB_531793 |
| Goat anti-Parvalbumin (IHC 1:2000) | Swant | Cat# PVG-213, RRID:AB_2650496 |
| Rabbit anti-Somatostatin (IHC 1:500) | Peninsula Laboratories | Cat# T-4103.0050, RRID:AB_518614 |
| Rabbit anti-TBR1 (WB 1:1000) | Abcam | Cat# ab31940, RRID:AB_2200219 |
| Rabbit anti-TBR1 (IHC 1:500) | Millipore | Cat# AB10554, RRID:AB_10806888 |
| Donkey anti-Rabbit IgG IRDye 800CW (WB 1:10,000) | LI-COR Biosciences | Cat# 926-32213, RRID:AB_621848 |
| Donkey anti-Mouse IgG (H+L), Alexa Fluor 488 (IHC 1:500) | Thermo Fisher Scientific | Cat# A-21202, RRID:AB_141607 |
| Donkey anti-Rabbit IgG (H+L), Alexa Fluor 546 (IHC 1:500) | Thermo Fisher Scientific | Cat# A10040, RRID:AB_2534016 |
| Donkey anti-Goat IgG (H+L), Alexa Fluor 647 (IHC 1:500) | Thermo Fisher Scientific | Cat# A-21447, RRID:AB_2535864 |
| Donkey anti-Rat IgG (H+L), Alexa Fluor 647 (IHC 1:500) | Jackson ImmunoResearch Labs | Cat# 712-605-153, RRID:AB_2340694 |
| **Chemicals, peptides, and recombinant proteins** | | |
| BtsCI | New England Biolabs | Cat# R0647S |
| BtsIMutI | New England Biolabs | Cat# R0664S |
| DiI Stain | Invitrogen | Cat# D3911 |
| **Critical commercial assays** | | |
| RNeasy Lipid Tissue Mini Kit | QIAGEN | Cat# 74804 |
| ProtoScript II First Strand cDNA Synthesis Kit | New England Biolabs | Cat# E6560S |
| PrimeTime Std qPCR Assay, *Tbr1* Exon Location 4–6, 6-FAM/ZEN/IBFQ | Integrated DNA Technologies | Mm.PT.58.42090842 |
| PrimeTime Std qPCR Assay, *Actb* Exon Location 5–6, SUN/ZEN/IBFQ | Integrated DNA Technologies | Mm.PT.39a.2221484 3.g |

| **Experimental models: Organisms/strains** | | |
|---|---|---|
| Mouse: $Tbr1^{KO}$ ($Tbr1^{tm1Jlr}$) | Mutant Mouse Resource & Research Centers | MMRRC:030263-UCD, RRID:MMRRC_030263-UCD |
| Mouse: $Tbr1^{A136PfsX80}$ | This study | N/A |
| Mouse: $Tbr1^{K228E}$ | This study | N/A |
| Mouse: $Tbr1^{\Delta 348–353}$ | This study | N/A |
| **Oligonucleotides** | | |
| CRISPR/Cas9 sgRNAs targeting *Tbr1*, see Table S1 | This study | N/A |
| CRISPR/Cas9 knock-in donors, see Table S1 | This study | N/A |
| Primers for genotyping, see Table S1 | This study | N/A |
| **Software and algorithms** | | |
| CFX Manager | Bio-Rad Laboratories | RRID:SCR_017251 |
| ImageJ2/FIJI | (Ruedin et al., 2017) | https://fiji.sc/; RRID:SCR_002285 |
| Image Studio | LI-COR Biosciences | RRID:SCR_015795 |
| Prism 9 | GraphPad Software | RRID:SCR_002798 |
| Sequencher | Gene Codes Corporation | RRID:SCR_001528 |
| ZEN Blue | Zeiss | RRID:SCR_013672 |
